# Wounding triggers invasive progression in human basal cell carcinoma

**DOI:** 10.1101/2024.05.31.596823

**Authors:** Laura Yerly, Massimo Andreatta, Josep Garnica, Charlée Nardin, Jeremy Di Domizio, François Aubin, Michel Gilliet, Santiago J. Carmona, François Kuonen

## Abstract

The interconnection between wound healing and cancer has long been recognized, as epitomized by the expression “cancer is a wound that does not heal”. However, the impact of inducing a wound, such as through biopsy collection, on the progression of established tumors remains largely unknown. In this study, we apply single-cell spatial transcriptomics to characterize the heterogeneity of human basal cell carcinoma (BCC) and identify a wound response gene program as the most prominent feature of highly invasive BCC. To explore the causal relationship between wounding and cancer invasive progression, we perform a longitudinal experiment to compare human tumors at baseline and one week post-biopsy. Our results demonstrate that biopsy collection triggers, in proximity of the wound, the same transcriptional cancer cell state observed in highly invasive BCC. This cancer cell transcriptional switch is coupled with morphological changes and the transcriptional reprogramming of cancer-associated fibroblasts (CAFs). This study provides evidence that wounding triggers invasive progression of established human tumors and warrants further research on the potentially harmful effects of biopsies and wound-inducing treatments.

## Introduction

Basal cell carcinoma (BCC), the most common cancer in humans, exhibits significant inter– and intra-tumoral morphological variability. Its histological patterns range along a spectrum from nodular to infiltrative forms^1,2^. The progression from nodular to infiltrative BCC is linked to increased local invasiveness and a higher risk of recurrence following primary treatments such as surgery or radiotherapy^3–5^. Relapsing, “advanced” BCCs, which are not eligible for surgery or radiotherapy, are typically treated with Hedgehog pathway inhibitors (HHI), targeting the primary oncogenic driver in BCC^6,7^. Considering that resistance to HHI occurs in up to 60% of treated patients^8^, investigating the relationship between nodular-to-infiltrative transition and treatment resistance could offer novel therapeutic insights and lead to more effective treatment strategies.

Previous studies have failed to identify somatic genomic alterations specific to infiltrative BCCs^9,10^, suggesting that invasive progression may primarily result from non-genetic mechanisms. Accordingly, prior research has shown that the transition from nodular to infiltrative morphology is a rapid and inducible process, accompanied by progressive transcriptional changes in cancer cells^11,12^. However, the mechanisms driving nodular-to-infiltrative transition in human tumors are largely unknown and finer characterization has been limited by the lack of high-resolution spatial molecular profiling technologies. Recent advances in spatial transcriptomics technologies with single-cell resolution have enabled the measurement of cancer cell-specific transcriptional programs *in situ* along with their interacting stromal and immune cells in human tumors *ex vivo*^13–15^. Here, we aimed to characterize the transcriptional heterogeneity of BCC cancer cells along the nodular-to-infiltrative transition, and its association with resistance to HHI.

In this work, we identified a wound response gene program in cancer cells – coupled with the transcriptional reprogramming of cancer-associated fibroblasts – as the most prominent feature of highly invasive BCC. Tackling the causal relation between induction of a wound and increased invasiveness in BCC, we designed a longitudinal experiment to compare the transcriptional landscape and morphology of tumors at baseline and after biopsy.

## Results

### Spatially-resolved cancer cell transcriptional programs in human basal cell carcinoma

Human BCC tumors display substantial intra-tumoral morphological heterogeneity (**Fig. 1A**). We hypothesized that this variability is reflected in the transcriptome by distinct patterns of activation of gene expression programs. To identify such programs, we performed an unsupervised meta-analysis of single-cell RNA-seq data from BCC tumors of 11 patients (**Fig. 1B** and **Fig. S1**; **Table S1**) (GSE181907^12^; E-MTAB-13085^16^). Using GeneNMF, a newly developed tool for multi-sample gene program discovery based on non-negative matrix factorization (see Methods), we identified seven recurrent gene programs (“meta-programs” or MPs) that consistently explain the cancer cell transcriptional heterogeneity across patients (**Fig. 1C**; **Table S2**). Comparison of these MPs with those identified in the comprehensive pan-cancer analysis by Gavish *et al.*^17^ revealed that MP1 and MP6 correspond to previously identified cell cycle and stress response MPs, respectively. The remaining MPs did not show significant association with previously reported cancer cell programs (**Fig. 1D** and Methods). Gene set enrichment analysis and manual examination of the MP genes suggested potential roles in cell radial organization for MP2 (*LAMB1* and *LRP8)*; in epidermal differentiation for MP3 (*CLDN4*, *CALML5*, *LGALS7B* and *KRT75*); in regulation of proliferation for MP4 (*SFRP5*, *ALK* and *CRNN*); in energy metabolism for MP5 (*MYL9* and *CSRP1*) and in epidermal response to wounding for MP7 (*TIMP1*, *IFITM3*, *KRT6A* and *INHBA*) (**Fig. 1D**; **Fig S2A**).

**Figure 1.**
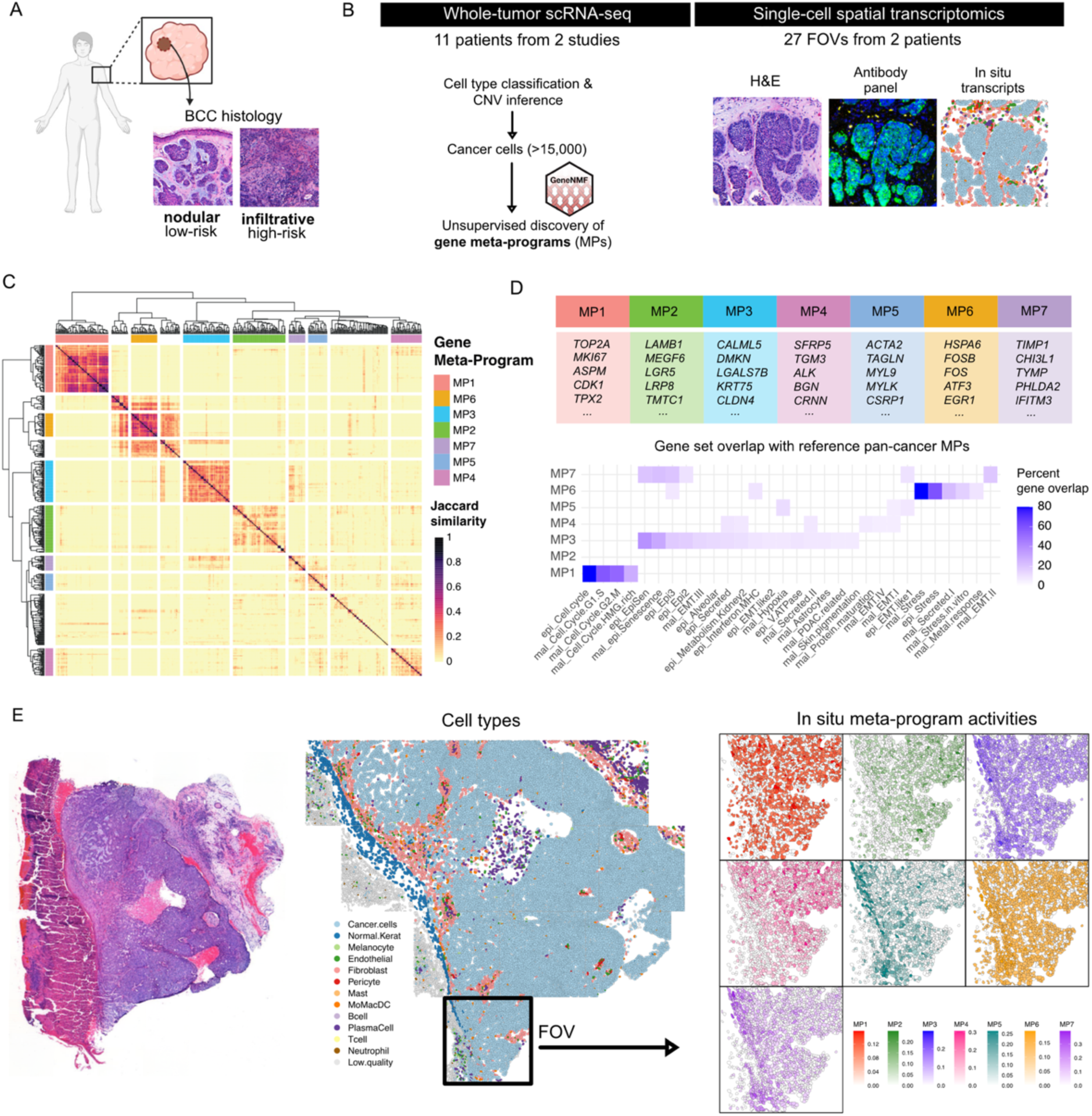
Spatially-resolved cancer cell transcriptional programs in human basal cell carcinoma. **A)** BCC displays histological heterogeneity that can be classified as nodular or infiltrative. Nodular BCC is characterized by nests of cancer cells with radial organization, separated from a loose myxoid stroma by a well-defined basement membrane. In contrast, infiltrative BCC displays branching strands of cancer cells embedded in a dense fibrotic stroma. **B)** Whole-tumor scRNA-seq was performed for 11 patients to study BCC transcriptional heterogeneity; cancer cells were computationally identified and subjected to gene meta-programs discovery using a new tool described in this paper, called GeneNMF (https://CRAN.R-project.org/package=GeneNMF). Single-cell spatial transcriptomics with CosMx SMI was performed to describe the spatial distribution of transcripts and their association with tumor morphology. **C)** Pairwise similarity (Jaccard index) between gene programs derived from individual samples, defining 7 clusters of robust and non-redundant programs, termed meta-programs (MP). **D)** Top 5 genes specific for each MP, and percent gene overlap between the 7 MPs and the reference pan-cancer meta-programs in malignant (mal) and normal epithelial (epi) cells defined by Gavish et *al*. (2023). **E)** Representative slice of human BCC analyzed with CosMx SMI; from left to right are shown H&E staining, predicted cell types from *in situ* transcripts, and activity of the seven meta-programs identified by scRNA-seq. FOV: Field-of-View.

To examine the spatial distribution of the identified MPs in the tumoral tissue, we performed single-cell spatial transcriptomics of human BCCs from 27 Fields-of-View (FOVs, 0.26 mm^2^-areas) distributed over 2 tumors, using CosMx SMI (Spatial Molecular Imager) technology with a 6’175 gene panel^18^ (**Fig. 1B**; **Fig S3**, **S4**). Using a supervised approach based on the expression profiles of scRNA-seq BCC data as reference^19^ (see Methods), we confidently classified single cell-resolved *in situ* transcriptomes into cancer cells, normal keratinocytes, endothelial cells, pericytes, fibroblasts, melanocytes, and immune cells, including T cells, B cells, plasma cells, mast cells, neutrophils, and mononuclear phagocytes (i.e. monocytes, macrophages, or dendritic cells) (**Fig. 1E**; **Fig. S5**). Activation of the identified MPs in cancer cells was consistently detected in *in situ* transcriptomes, and followed a similar cell-to-cell correlation pattern as in scRNA-seq data (**Fig. 1E**; **Fig. S2B-C**). Altogether, we identified 7 MPs explaining the transcriptional heterogeneity of human BCC, and their spatial distribution in the tissue at single-cell resolution.

### Cancer cell-intrinsic transcriptional programs underpin invasive patterns

BCC histologic morphology ranges from nodular to infiltrative patterns, with progression marked by increased invasiveness –a defining hallmark of cancer^20^. To assess whether morphological changes are associated with differential activation of transcriptional programs, we took advantage of the 2 human BCCs analyzed by single-cell spatial transcriptomics displaying mixed morphology – *i.e.* containing both nodular and infiltrative areas (**Fig. 2A; Fig. S4**). Individual tumor areas (*i.e.* FOVs) were assessed for global morphology, cell type composition and, within cancer cells, homotypic versus heterotypic physical cell-cell interactions (see Methods). As expected, FOVs with infiltrative morphology (as determined by H&E histological examination) displayed higher morphological complexity as measured by the ratio of heterotypic to homotypic cancer cell interactions (**Fig. 2B**) and perimeter/area ratio (**Fig. S6**), when compared to FOVs with nodular morphology. Interestingly, infiltrative tumor regions displayed a lower proportion of cancer cells (**Fig. 2C**), mirrored by a statistically significant increase in all detected immune cell types as well as in fibroblasts and endothelial cells, but not in pericytes or melanocytes (**Fig. 2D**). This indicates that BCC invasiveness is associated with local inflammation and peritumoral tissue remodeling, as previously observed^2,11^.

**Figure 2.**
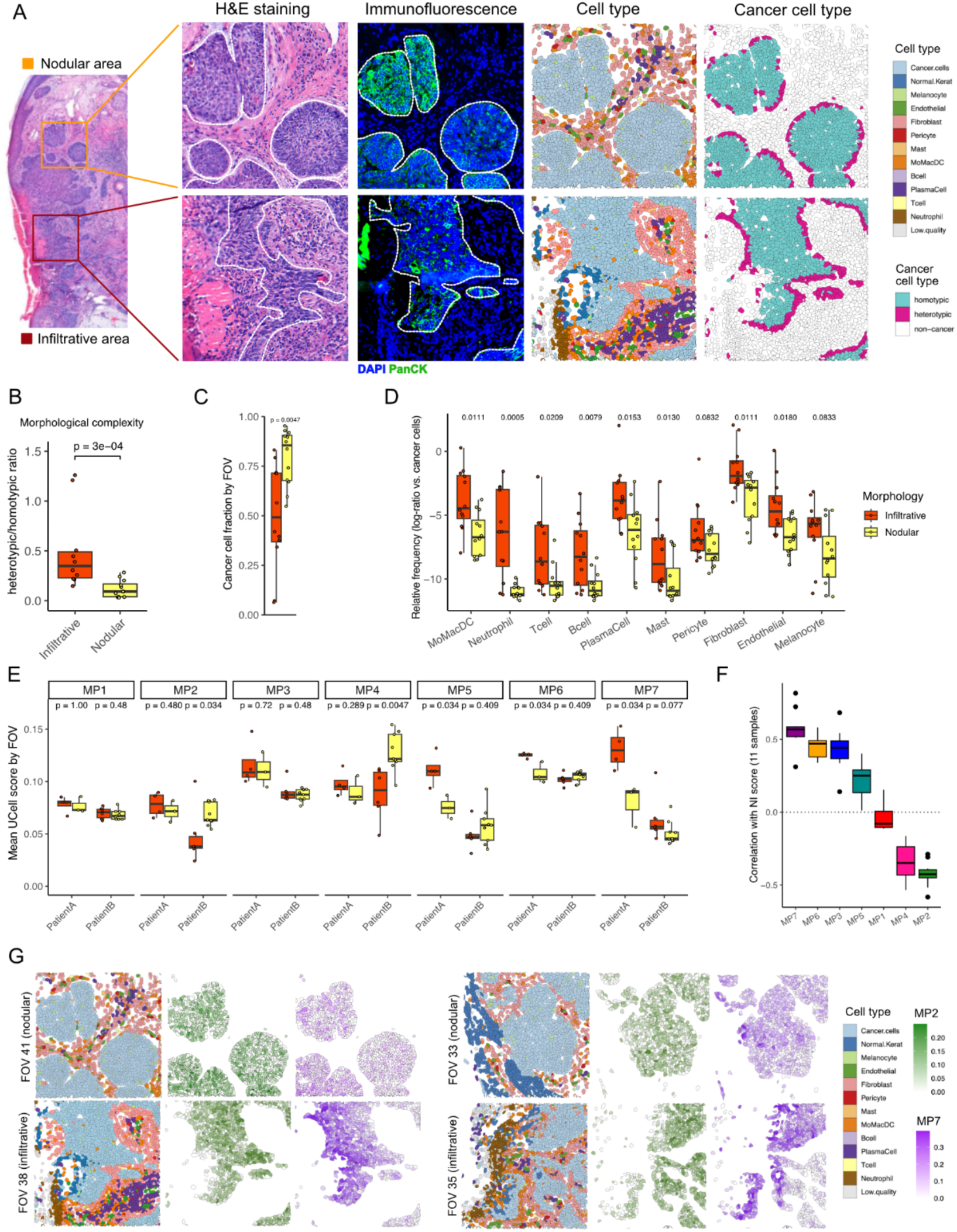
Cancer cell-intrinsic transcriptional programs underpin invasion patterns in human basal cell carcinoma. **A)** Representative Fields-of-View (FOVs) from nodular and infiltrative BCC areas. From left to right are shown H&E staining, immunofluorescence for DAPI and panCK, cell type annotation from spatial *in situ* transcripts (CosMx SMI), and homotypic vs. heterotypic cell-cell interactions in cancer cells. H&E and spatial transcriptomics are derived from consecutive slides. **B)** Morphological tumor complexity as measured per CosMx FOV as measured by the ratio of heterotypic/homotypic cancer cell interactions. **C)** Cancer cell fraction in nodular vs. infiltrative FOVs. **D)** Relative frequency of immune and stromal cell types (measured as the log-ratio relative to cancer cells) in nodular vs. infiltrative FOVs. **E)** Meta-program activity (UCell score) for MP1-MP7, in FOVs from two different patients. **F)** Correlation of MP activity with nodular-to-infiltrative score (NI score) in 11 scRNA-seq BCC samples. **G)** Representative FOVs for two nodular and two infiltrative tumor areas, highlighting cell type composition and spatial distribution of meta-programs MP2 and MP7. p-values: Wilcoxon rank-sum test.

Interestingly, we found that several meta-programs were differentially activated in nodular and infiltrative FOVs. A consistent association with the infiltrative pattern was found for MP7 – and to a lesser extent for MP5 and MP6. In contrast, MP2 and MP4 activation appeared to be associated with the nodular pattern (**Fig. 2E**). To support these observations, we leveraged previous work in which we applied GeoMx spatial transcriptomics to identify gene signatures characteristic of highly-infiltrative and highly-nodular BCC tumor niches^12^. These signatures allowed us to calculate a nodular-to-infiltrative (NI) score for each single cancer cell, ranging from –1 in fully nodular to 1 in fully infiltrative BCC cancer cells (see Methods). We thus examined whether the activation pattern of our MPs was associated, in our scRNA-seq cohort (**Fig. 1B**), with the nodular or infiltrative pattern as quantified by the NI score. In agreement with the single-cell spatial transcriptomic data, MP2 and MP7 displayed the strongest association with the nodular and the infiltrative pattern, respectively (**Fig. 2F**). These correlations were consistent across individual patients (**Fig. S7A-B**). Further analysis of MP activity by spatial transcriptomics confirmed that MP2 and MP7 (for FISH assays represented by markers *MPPED1* and *KRT6A*, respectively, see **Fig. S8A**) were negatively correlated and their activation reflected morphological complexity (**Fig. 2G**; **Fig. S8B**, **Fig. S7C**).

In sum, we identified cancer cell-intrinsic gene expression programs underlying BCC morphological patterns and identified a gene meta-programs axis (MP2 Η MP7), coupled with TME changes, that is strongly associated with BCC invasive progression.

### Invasive progression is associated with resistance to Hedgehog pathway inhibitors

To characterize the axes of MP activation in increasingly aggressive skin tumors, we analyzed publicly available scRNA-seq data from advanced, HHI-resistant BCCs^21^ as well as cutaneous squamous cell carcinoma (cSCC)^22^, a more aggressive, keratinocyte-derived cancer type^23^. Notably, in both cohorts cancer cells displayed lower MP2 and increased MP7 activation compared to nodular or infiltrative treatment-naive BCCs (**Fig. 3A**, **Fig. S9**), suggesting that invasive progression parallels the development of HHI resistance. To confirm this, we assembled a new cohort of patients with advanced – i.e. eligible for HHI therapy – BCCs (**Fig. S10A**). Compared to early BCCs or advanced naive BCCs (irrespective of nodular or infiltrative dominant morphology), HHI-resistant BCCs displayed a significantly higher morphological complexity (**Fig. S10B-C**), indicating increased invasiveness. Furthermore, in two HHI-resistant patients for which pre– and post-treatment biopsies were available, we found increased morphological complexity and MP7/MP2 ratio upon HHI treatment relapse (**Fig. 3B**; **Fig. S10D**).

**Figure 3.**
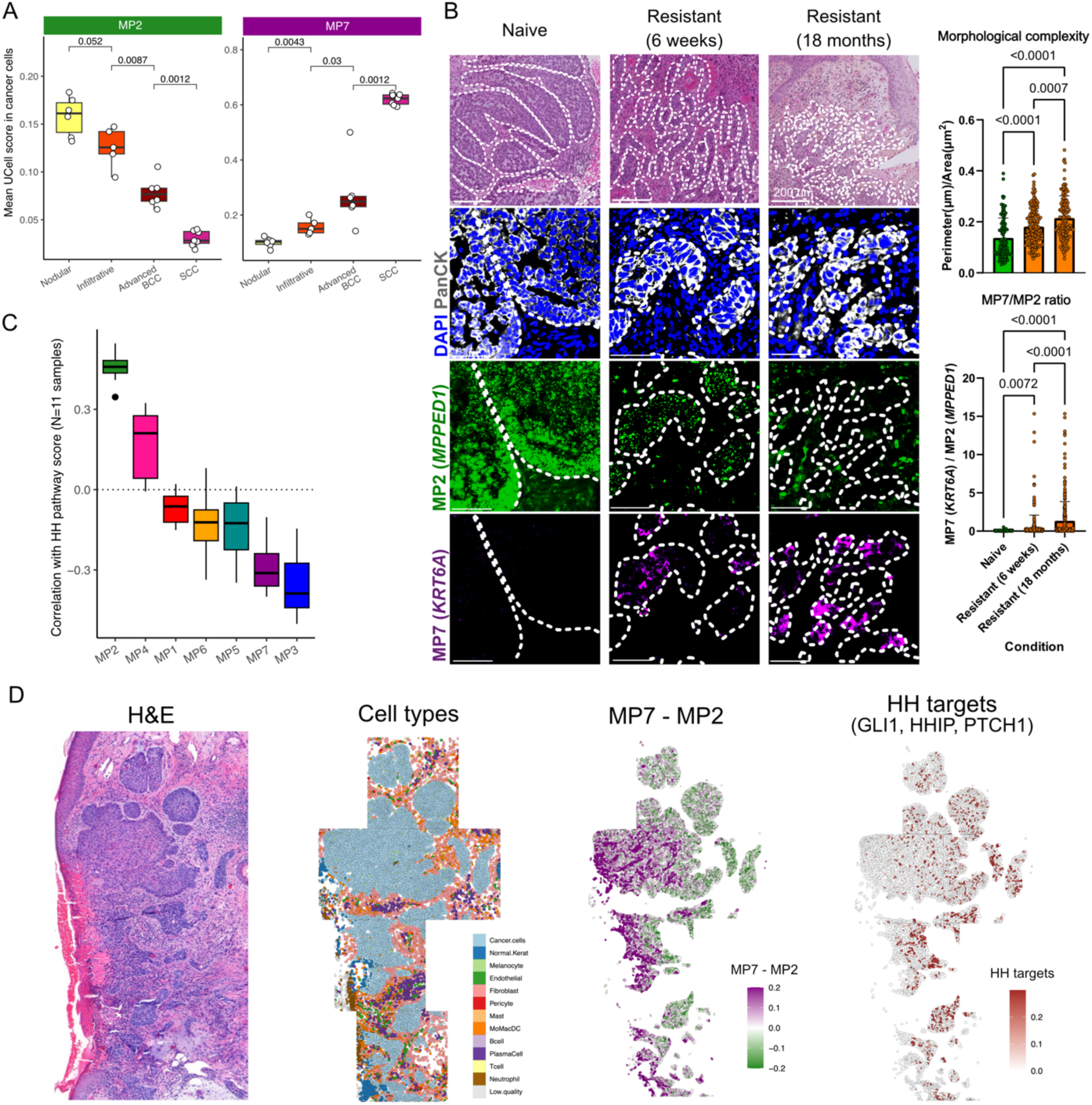
Invasive progression is associated with resistance to hedgehog pathway inhibitors. **A)** Mean signature score per sample for meta-programs MP2 and MP7, in cancer cells from scRNA-seq samples of nodular BCCs, infiltrative BCCs, advanced therapy-resistant BCCs, and cutaneous squamous cell carcinomas (cSCCs). **B)** Patient-matched treatment-naive and treatment-resistant (after 6 weeks and 18 months treatment) samples shown for representative H&E and immunofluorescent images; morphological complexity (measured by perimeter/area ratio; top right panel); and MP7/MP2 ratio (assessed by the ratio of mean pixel intensity of MP7 marker *KRT6A* over mean pixel intensity of MP2 marker *MPPED1*; bottom right panel). P-values: One-way ANOVA and Tukey’s multiple comparison test**. C)** Correlation of MP activity with KEGG hedgehog pathway score in 11 scRNA-seq BCC samples. **D)** Spatial distribution of MP7 minus MP2 signature score, and HH targets (*GLI1*, *HHIP*, *PTCH1*) signature score in a representative tissue area analyzed with CosMx SMI *in situ* transcriptomics.

Phenotypic plasticity is a hallmark of cancer resistance to targeted therapy^20,24^ and is largely implicated in shifting advanced BCC away from dependency on the HH pathway^25–28^. To characterize the relation between HH pathway and MPs activation – regardless of HHI treatment, we analyzed scRNA-seq data of early, naive BCC. MP2 was the meta-program most strongly positively correlated with HH pathway activity, while MP7 displayed a strong negative correlation (**Fig. 3C**). These correlations were consistent across individual patients (**Fig. S11**). Further analysis of the spatial distribution of MPs and HH pathway activation using spatial transcriptomics confirmed the progressive shutdown of the HH pathway along the MP2 to MP7 axis (**Fig. 3D; Fig. S12**).

Altogether, we identified a transcriptional shift in cancer cells, which couples local invasiveness with reduced HH signaling. Such reprogramming might foster resistance to HH-targeting therapy by shifting cancer cells away from HH-dependency in advanced BCCs.

### Tumor wounding triggers invasive progression in human BCC

MP7 genes have a presumed role in epidermal wound healing, while antagonist MP2 includes *LGR5*, a marker of hair follicle stem cells – which have rapid adaptive capacity during wound healing^29^. This suggests that the nodular-to-infiltrative transition (and related MP2 to MP7 transcriptional axis) might reflect tumor response to wounding. To investigate the association of tumor wounding with invasive progression in BCC tumors, we first turned to a typical setting of skin tumor injury – namely ulceration^1,30,31^. We extracted BCC samples classified as nodular-ulcerated based on histopathological reports from our tissue biobank (see Methods). Remarkably, in nearly all samples, ulcerated areas displayed increased invasiveness, reflected by higher morphological complexity (**Fig. S13A-B**), downregulated expression of *MPPED1* (a MP2 marker) and upregulated expression of *KRT6A* (a MP7 marker) in cancer cells (**Fig. S13C-D**). These results establish a link between wounding and the MP2-MP7 transcriptional switch in cancer cells. While this association might result from highly invasive BCC tumors outgrowing their blood supply and leading to ulceration, it is an intriguing possibility that wounding might be a factor promoting tumor invasive capacity.

To test whether wounding could trigger nodular-to-infiltrative transition in human BCC, we designed a longitudinal experiment to compare tumors taken before and after inducing a wound by punch biopsy. We selected four patients with non-ulcerated, non-biopsied, nodular BCC tumors. For each patient, we collected three samples: *i)* from baseline (first, diagnostic biopsy), and 1 week later *ii)* from the previously biopsied, “wounded” site, and *iii)* from a distant, “unwounded” site in the same tumor (**Fig. 4A**). All samples were subjected to H&E staining and, for two patients, to spatial transcriptomics using CosMx SMI technology (**Fig. S14-15**) and/or scRNA-seq. Individual CosMx FOVs from wounded samples were further annotated according to their proximity to the wound (into Wound-Far and Wound-Close, **Fig. S15**). Notably, we found that one week after punch biopsy-induced wounding, the wounded tumor sites displayed a transition from a nodular to an infiltrative morphology, as reflected by increased morphological complexity measured on H&E (**Fig. 4B**; **Fig. S16A-B**). This observation was confirmed on the CosMx spatial transcriptomics data, by measuring the ratio of heterotypic/homotypic cancer cells interactions within individual FOVs (**Fig. 4C**). In terms of meta-program activity, MP7 (and to a lesser extent MP3 and MP5) displayed a significant increase in wounded samples compared to the baseline and unwounded tumors. Conversely, MP2 was significantly downregulated in proximity of the wound compared to baseline (**Fig. 4D**). A similar trend was observed in both patients processed by scRNA-seq (**Fig. S16C**). Remarkably, MP2 and MP7 marker genes were among the most differentially expressed genes in the wounded vs. baseline or unwounded conditions (**Fig. 4E** and **Fig. S16D**), indicating that these gene programs captured the most prominent transcriptional changes that are induced upon tumor wounding.

**Figure 4.**
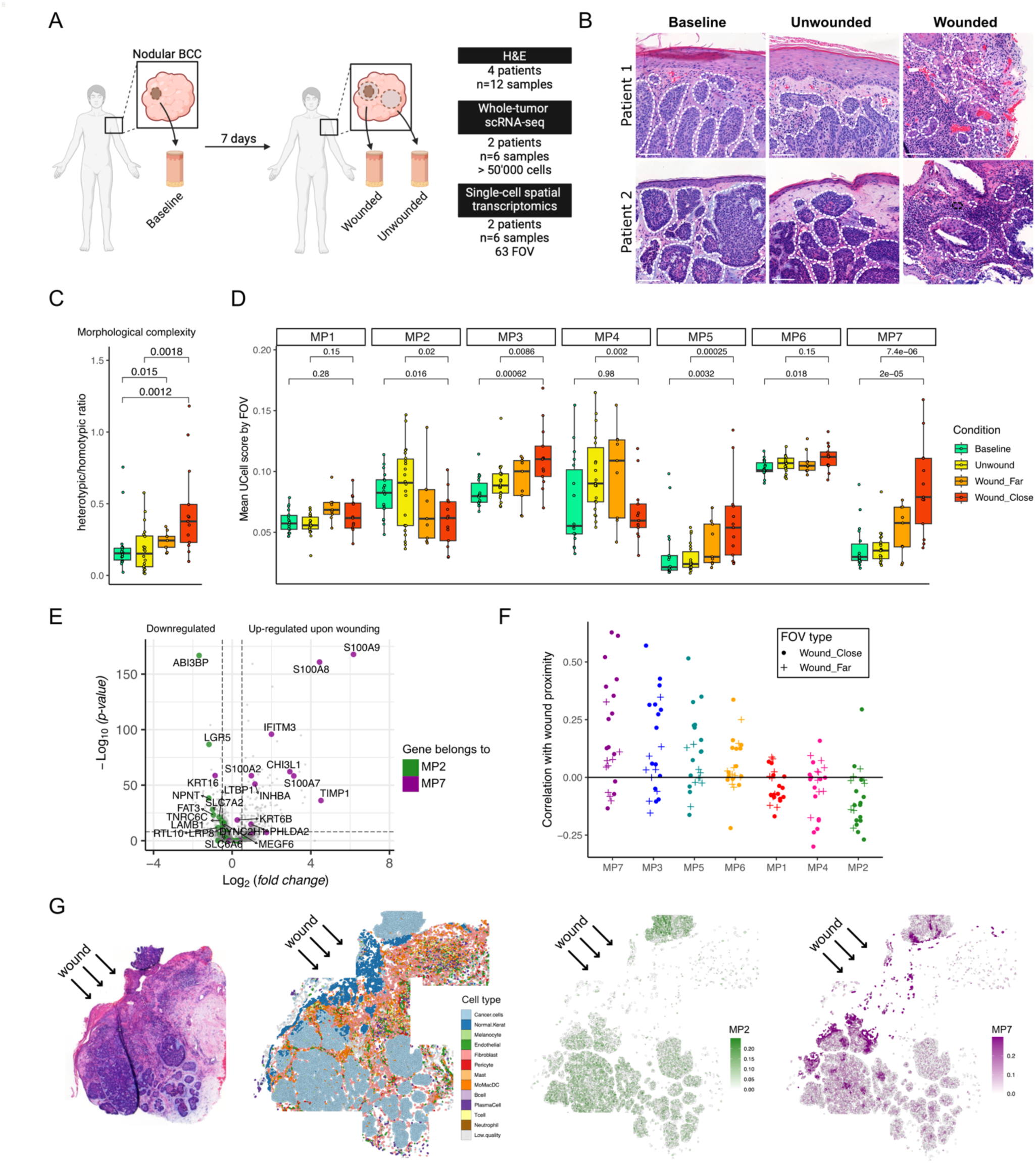
Tumor wounding triggers invasive progression in human BCC. **A)** Experimental design: for each patient, samples were collected with a first biopsy (Baseline) and 1 week later from the previously biopsied site (Wounded) and at a distant site (Unwounded) in the same tumor. Samples were processed for H&E, whole-tumor scRNAseq and single-cell spatial transcriptomics with CosMx SMI. **B)** Representative scan of H&E images from 2 BCC patients at baseline and 7 days later in the wounded and unwounded areas. White dotted lines depict the basement membrane zone (BMZ). Scale bars indicate 100 µm. **C)** Morphological tumor complexity per CosMx FOV as measured by the ratio of heterotypic/homotypic cancer cells, in the indicated sample types. **D)** Meta-program activity (UCell score) for MP1-MP7 in *in situ* transcripts from CosMx SMI FOVs. **E)** Differentially expressed genes between Wound vs. Baseline/Unwound samples for a representative patient (data for second patient in Figure S16). Genes contained in MP2 and MP7 are highlighted green and purple respectively. **F)** Correlation coefficient between MP activity and proximity to the wound; each point represents correlation within one FOV, MPs are sorted by decreasing average correlation. **G)** Tiled FOVs for one representative wounded tumor slide, with approximate wound site marked by arrows; from left to right are shown H&E staining, predicted cell types from *in situ* transcripts, and spatial meta-program activity for MP2 and MP7. p-values: Wilcoxon rank-sum test.

We next evaluated how MP activation related to the wound proximity (**Fig. S14-15**). We calculated, in each FOV, the relative distance of each cancer cell to the wound and examined MP activation as a function of this distance. Remarkably, MP7 activity showed the strongest correlation with wound proximity, while MP2 activity progressively increased in cancer cells further from the wounded site (**Fig. 4F**). Such correlations were globally stronger when considering FOVs close to wounding (Wound_Close) compared to FOVs far from wounding (Wound_Far) (**Fig. 4F**). This directionality of MP2 and MP7 activity toward the wounding site further supports that the activation of these gene programs is orchestrated by tumor wounding (**Fig. 4G**).

Altogether, our data show that injury locally triggers a cancer cell switch from a non-invasive to an invasive state. This observation supports the role of wounding as a driver of transcriptional reprogramming and invasive progression in human BCC.

### Wounding induces a cancer-associated fibroblast state characteristic of invasive and advanced BCCs

Our results revealed a transcriptional switch and spatial reorganization of cancer cells that is characteristic of both invasive progression and acute wound response in human BCC. To investigate whether the interconnection between these two processes extends beyond cancer cells, we examined the changes induced by wounding on the TME.

Based on our spatial transcriptomics data, we observed that the proportion of cancer cells was reduced upon wounding (**Fig. 5A**), in favor of increased infiltration of all immune and most stromal cell types (**Fig. 5B-C**) – as also previously observed in infiltrative BCCs (**Fig. 2D**). Using graph-based spatial network analysis, we analyzed the spatial relationships between specific cell types and how they change upon wounding (**Fig. 5C**). Interestingly, wounding not only increased significantly the proportion of T cells and monocytes/macrophages/dendritic cells, but these cell types also tended to co-localize with each other and with fibroblasts. As expected, melanocytes were mostly found embedded within the tumor mass and were therefore closely linked spatially to cancer cells. Neutrophils were only observed in detectable numbers in FOVs close to the wound and did not appear to consistently co-localize with other cell types (**Fig. 5C**). To confirm our observations using a different assay, we analyzed the stromal and immune cell types in scRNA-seq of baseline, unwounded and wounded samples for two patients. We confirmed that wounded samples displayed a ∼10-fold increase in the relative abundance of immune cells (including T cells, monocytes, macrophages and dendritic cells) and cancer-associated fibroblasts (CAFs), as well as a modest increase of endothelial cells and a decrease of pericytes (**Fig. S17A**). The strongest transcriptional changes (in terms of number of differentially expressed genes) upon wounding were captured in dendritic cells, cancer-associated fibroblasts (CAF), monocyte/macrophages (MoMac) and pericytes (**Fig. 5D**).

**Figure 5.**
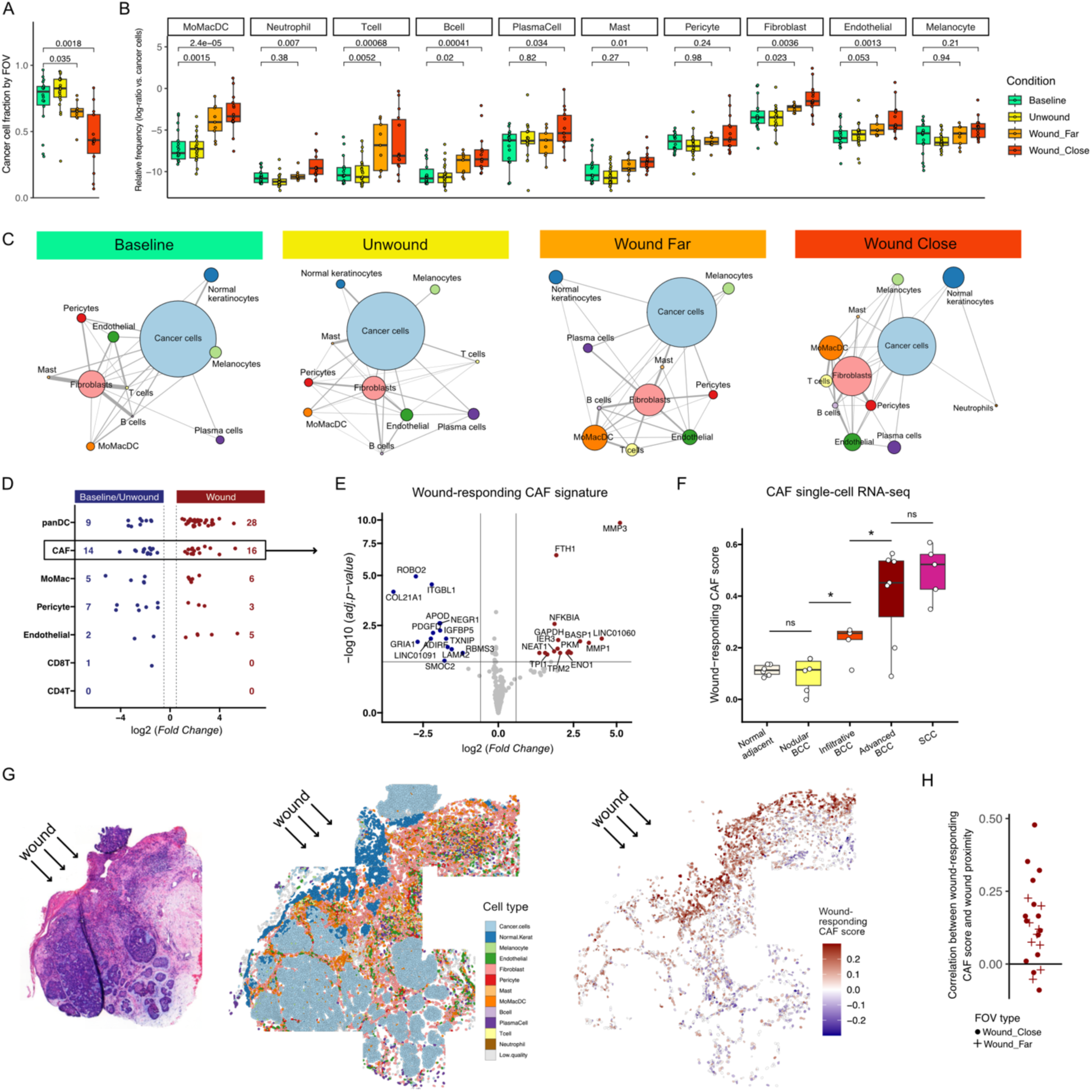
Wounding induces a cancer-associated fibroblast state characteristic of invasive and advanced BCCs. **A)** Fraction of cancer cells in CosMx SMI Fields-of-View (FOVs) for the four indicated conditions. **B)** Relative frequency (calculated as log-ratio with respect to cancer cells) of immune and stromal cell types; each point represents one FOV. **C)** Cell type interactome in the four indicated conditions; node size is proportional to the cell type frequency, edge length reflects average distances between cell types in a physical model of springs (Kamada-Kawai algorithm); edge width is proportional to the inverse of the standard deviation of cell type distances. **D)** Absolute number and log2 fold-change of differentially expressed genes (DEGs) between Wound and Baseline/Unwound scRNA-seq samples, for each of the indicated cell types. **E)** Volcano plot of DEGs for cancer-associated fibroblasts (CAFs); genes with p_adj < 0.05 and log2 (fold-change) < 0.5 were used to build a “wound-responding CAF” signature. **F)** Mean “wound-responding CAF” score per sample, in fibroblasts from scRNA-seq samples of normal adjacent skin, nodular BCCs, infiltrative BCCs, advanced therapy-resistant BCCs, and cutaneous squamous cell carcinomas (cSCCs). **G)** Tiled FOVs for one representative wounded tumor slide, with approximate wound site marked by arrows; from left to right are shown H&E staining, predicted cell types from *in situ* transcripts, and spatial distribution of “wound-responding CAF” signature score. **H)** Correlation coefficient between “wound-responding CAF” signature score and proximity to the wound; each point represents correlation within one FOV. p-values: Wilcoxon rank-sum test.

We next asked whether these acute wound-induced TME cell states might be present in chronically infiltrative BCCs, similarly to our previous findings for MP7 in cancer cells. While we found no association for the immune cells states, the differentially-expressed genes in CAFs upon wounding were collectively upregulated in infiltrative vs. nodular BCCs (**Fig. 5E**, **Fig. S17B**). These included several genes with a presumed role in the fibroblast response to wound healing in healthy skin (*MMP3*, *FTH1*, *IER3*^32^) (**Table S3**). The wound-responding CAF gene program was not only significantly more active in CAFs in infiltrative BCCs compared to those in nodular or normal adjacent samples (**Fig. 5F**), but was also increased in proximity to the wound (**Fig. 5G-H**, **Fig. S17C**). The pattern was further confirmed on the GeoMx data, where the CAF program activity was higher in infiltrative vs. nodular BCC stroma-rich regions (**Fig. S17D**). These results indicate that –similarly to MP7 in cancer cells– the wound-responding gene program in CAFs is strongly upregulated both acutely upon tumor wounding and chronically in highly infiltrative BCCs. Strikingly, the wound-responding CAF gene program displayed highest activity in human samples of advanced, HHI-resistant BCC^21^ and cSCC^22^ (**Fig. 5F**) – aggressive skin tumor types which were also shown to downregulate MP2 and upregulate MP7 (**Fig. 3A**).

Altogether, these data support the notion that infiltrative progression triggered by tumor wounding is accompanied by changes in the TME, in particular by quantitative and qualitative changes in the CAF population. Additionally, CAF reprogramming acutely induced upon biopsy largely overlaps with the differences observed between non-invasive and invasive BCC tumors (further exacerbated in advanced BCCs), suggesting that the invasive alterations triggered by induced wounding may persist over time.

## Discussion

Multiple cellular and molecular processes involved in the wound healing response are often also active and dysregulated in tumors^20,33^. While the tumor-promoting effect of wound-induced inflammation on early stages of tumor formation has been well documented^34–38^, it is unclear how wounding can impact cancer cell progression in established tumors.

In this study, we demonstrated that wounding –such as that caused by biopsy collection– in BCC patients triggers a transcriptional shift in cancer cells, which transition from a non-invasive to an invasive state. One week after biopsy of non-invasive nodular BCCs, in the wound proximity, cancer cells and cancer-associated fibroblasts (CAFs) display a strong transcriptional reprogramming pattern that we found to be characteristic of high-risk infiltrative BCC and advanced BCC, as well as of more aggressive keratinocyte-derived cSCCs. The reprogramming of cancer cells and CAFs may be initiated by the inflammatory response, which is rapidly activated following tissue injury^39–41^. This is supported by the expression of acute inflammatory response genes (e.g. *IFITM3* and *CHI3L1* in cancer cells, *NFKB1A* and *TNFAIP6* in CAFs) one week upon wounding in tumor areas in close proximity to the wound. *CHI3L1*, a gene of cancer cell metaprogram MP7, has also recently been identified as a marker of an inflammation-enriched tumor state in a mouse model of BCC^42^.

An outstanding important question is how the acute wound response in tumors persists over time. Ulcerated tumors typically fail to heal^43^, perpetuating a persistent inflammatory signal that potentially sustains tumor wound response in the vicinity of chronic ulceration. Interestingly, our data show that, unlike wound-induced transcriptional changes in immune cells, wound-responding cancer cell and CAF states persist in infiltrative tumors. One possible explanation is that pathophysiological processes associated with wound healing other than inflammation, such as hypoxia^44^, are at play. Alternatively, wound-responding cancer cell and CAF states may sustain themselves without requiring persistent inflammatory signals. Wound-responding CAFs, by remodeling the extracellular matrix (ECM), may dampen inflammation while supporting cancer cell progression in a self-sustained positive feedback loop, like keratinocytes and dermal fibroblasts during wound healing^45,46^. A deeper study of the molecular crosstalk induced upon injury between cancer cells and CAF – and maintained in infiltrative and advanced BCC – is thus expected to provide attractive therapeutic targets to revert wound-induced phenotypic plasticity and associated cancer progression.

MP7 includes genes implicated in epithelial cell migration (e.g. *TNC*, *TIMP1*, *ITGAV, ITGB6*) and epidermal barrier restoration (e.g. *KRT16*, S*100A7*), reflecting the specific organization of the wound migrating edge^47–50^. Interestingly, *ITGB6* (Integrin Subunit Beta 6) and *ITGAV* (Integrin alpha-V) code for the fibronectin receptor αVβ6, which mediates migration of keratinocytes to newly-formed, fibrotic stroma during wound healing^51,52^, and has been previously shown to mediate invasion in BCC^53^. Fibronectin is a major peritumoral constituent of infiltrative BCCs^11^ and its expression is directly regulated by Activin A signaling in both healthy skin fibroblasts and CAFs^54,55^. Upon wounding, we observe a rapid induction of *INHBA* in cancer cells. *INHBA* encodes Activin A and is persistently expressed in infiltrative BCCs^12^, indicating that wound-responding cancer cells may initiate CAF reprogramming towards ECM-remodeling features and thus provide a provisional matrix for cancer cell invasion^56,57^. While the parallel between dermal fibroblasts and CAFs support the similarity between wound healing and cancer, these results also raise the question of why in cancer, unlike in normal tissue, the wound healing process remains incomplete. Further insights may come from the investigation of discrepancies –rather than commonalities– between cancer and the late-stage of tissue repair.

Beyond the identification of tumor ulceration as a potential clinical indicator of an infiltrative BCC, our data raise questions about the risk-benefit balance of wounding procedures in clinical practice. Biopsies, the mainstay for diagnosing and grading cancers, as well as physical (surgery, radiotherapy, cryotherapy) or chemical (chemotherapy, photodynamic therapy) therapies induce tissue injury. Our data support a potential deleterious effect of these procedures as they may trigger local progression of residual cancer cells. This observation might explain the noted disparity between histological evaluations of biopsy samples and subsequent excisions in various solid epithelial cancers including BCC^58–64^, as well as the increased invasiveness of recurring BCCs after wound-inducing treatments^65,66^. Due to ethical considerations preventing long delays between diagnosis and treatment, our study did not address the long-term effects of tumor wounding.

Another important consideration is whether therapies that induce wounds contribute to resistance in residual cancer cells against subsequent lines of treatment, as suggested by our findings and prior clinical studies^67^. This highlights the need for additional therapeutic strategies to overcome cumulative resistance in cancer. In this context, stromal components may represent more attractive therapeutic targets compared to cancer cells, which are often characterized by genomic instability and cellular plasticity, resulting in therapy-resistant subclones^24^. CAF are highly responsive to various treatment regimens including chemotherapy and radiotherapy^68,69^, adopting a pro-fibrotic state linked to poor clinical outcome^70–72^. Consistently, we identify CAFs as a critical component of the tumor response to wounding, characterized by ECM-remodeling features linked to HHI resistance. As such, they emerge as a promising therapeutic target to overcome cumulative resistance. For instance, a recent study showed that targeting ECM-remodeling CAFs, which support glioblastoma cancer cell survival, restore sensitivity to standard-of-care therapy^73^.

Overall, our study describes profound histological, compositional and transcriptional changes induced by biopsy collection in human BCC tumors, supporting an association between wounding and cancer progression, and pinpointing wound-responding CAFs as attractive targets for otherwise therapy-resistant BCC.

### Study limitations

Due to ethical considerations preventing long delays between diagnosis and treatment, our study does not cover the full dynamics of alterations induced upon wounding, especially their long-term maintenance. Thus, the evidence supporting long-term consequences of the cancer cells transcriptional reprogramming induced by tumor wounding are indirect. Second, the deconvolution of cancer cell heterogeneity is based on a limited cohort of naïve BCC patients eligible for surgical treatment, and, as such, may not fully capture the heterogeneity of advanced BCCs. Finally, the association between wound-induced tumor changes and poorer prognosis (i.e. increased risk of recurrence or HHI therapy resistance) in the present study is based on observed correlations and would require clinical studies to establish a direct causal relationship.

## Acknowledgments

This work was supported by the SNSF fellowship PZ00P3-185926 (F.K.), PROMEDICA Stiftung (F.K.), and SNF fellowship 180010 (S.J.C). The authors would like to thank Dr. Daniele Tavernari (EPFL, UNIL) for helpful discussions on non-negative matrix factorization for gene program identification. We thank the Genomic Technologies Facility (GTF) for the scRNA-seq library preparation and sequencing. F.K. and M.G. are members of the SKINTEGRITY.CH collaborative research consortium.

## Author Contributions

L.Y., M. A., S.J.C. and F.K. conceived and designed the study and analyzed the experiments. L.Y. performed the wet-lab experiments. L.Y., M. A., J.G. and S.J.C. performed the computational analyses. L.Y., M. A., S.J.C. and F.K. wrote the manuscript with comments from co-authors. N.C., J.D.D, F. A. and M.G. provided access to the tissue biobank.

## Supplemental information

### Supplemental Tables

**Table S1:** Metadata table summarizing patient’s characteristics of samples used for scRNAseq and Cosmx analysis.

| ScRNA-seq | Cosmx | Sample | Age | Gender | Location |
| --- | --- | --- | --- | --- | --- |
| GSE181907 |  | BCC1 | 82 | M | left ear |
| GSE181907 |  | BCC2 | 78 | M | forehead |
| GSE181907 |  | BCC3 | 79 | M | left cheek |
| GSE181907 |  | BCC4 | 50 | F | nose |
| GSE181907 |  | BCC5 | 69 | F | nose |
| GSE181907 |  | BCC6 | 86 | M | ear |
| GSE181907 |  | BCC7 | 85 | M | upper lip |
| E-MTAB-13085 |  | BCC8 | 39 | M | cheek |
| E-MTAB-13085 |  | BCC9 | 73 | M | forehead |
| E-MTAB-13085 |  | BCC10 | 51 | M | temple |
| E-MTAB-13085 |  | BCC11 | 59 | M | temple |
| E-MTAB-13085 |  | BCC12 | 69 | F | ear |
| E-MTAB-13085 |  | BCC13 | 85 | M | cheek |
| E-MTAB-13085 |  | BCC14 | 70 | M | ear |
| E-MTAB-13085 |  | BCC15 | 91 | M | nose |
| GSE266665 |  | Patient 1 Baseline | 85 | M | neck |
| GSE266665 |  | Patient 1 Unwound | 85 | M | neck |
| GSE266665 |  | Patient 1 Wound | 85 | M | neck |
| GSE266665 | 10.5281/zenodo.14330690 | Patient 2 (scRNAseq)/<br>Patient D (Cosmx)<br>Baseline | 97 | F | cheek |
| GSE266665 | 10.5281/zenodo.14330690 | Patient 2 (scRNAseq)/<br>Patient D (Cosmx)<br>Unwound | 97 | F | cheek |
| GSE266665 | 10.5281/zenodo.14330690 | Patient 2 (scRNAseq)/<br>Patient D (Cosmx)<br>Wound | 97 | F | cheek |
|  | 10.5281/zenodo.14330690 | Patient A | 77 | M | temple |
|  | 10.5281/zenodo.14330690 | Patient B | 86 | M | neck |
|  | 10.5281/zenodo.14330690 | Patient C Baseline | 76 | M | cheek |
|  | 10.5281/zenodo.14330690 | Patient C Unwound | 76 | M | cheek |
|  | 10.5281/zenodo.14330690 | Patient C Wound | 76 | M | cheek |

**Table S2**. BCC meta-program genesets (as .xlsx file).

**Table S3**. Wound-responding CAF signatures (as .xlsx file).

**Table S4**. Meta-data for FOVs of CosMx SMI experiment (as .xlsx file).

### Supplemental Figures

**Figure S1. Identification of cancer cells from scRNA-seq of two human BCC cohorts. A)** UMAP embeddings by patient and by cell type annotation from the Yerly et al. cohort. Broad annotation was obtained by applying the scGate automated tool and refined by clustering analysis. Identification of cancer cells was performed using three criteria (patient mixing, signature scoring, and copy-number variation), outlined respectively in panels B, C and D. **B)** Local Inverse Simpson Index (LISI) by cell type, quantifying the number of patients having cells in any given neighborhood of cells. Cancer cells are expected to have more patient-specific transcriptomes than other cell types, and primarily cluster together with cells from the same patient. **C)** Scaled average expression for marker genes of major TME cell types, and UCell score for a BCC signature defined by Yerly et al. (2022). **D)** Copy-number variation (CNV) profile for a representative patient, confirming that normal keratinocytes and cancer cells primarily cluster together in terms of inferred CNVs. **E)** UMAP embeddings by patient and by cell type annotation from the Ganier et al. cohort. Broad annotation was obtained by applying the scGate automated tool and refined by clustering analysis. Identification of cancer cells was performed using three criteria (patient mixing, signature scoring, and copy-number variation), outlined respectively in panels F, G and H. **F)** Local Inverse Simpson Index (LISI) by cell type, quantifying the number of patients having cells in any given neighborhood of cells. Cancer cells are expected to have more patient-specific transcriptomes than other cell types, and primarily cluster together with cells from the same patient. **G)** Scaled average expression for marker genes of major TME cell types, and UCell score for a BCC signature defined by Yerly et al. (2022). **H)** Copy-number variation (CNV) profile for a representative patient, confirming that normal keratinocytes and cancer cells primarily cluster together in terms of inferred CNVs.

**Figure S2. Characterization of the meta-programs and their interrelation. A)** Gene set enrichment analysis (GSEA) between BCC meta-programs and Gene Ontology (GO) gene sets from MSigDB. Dot size is proportional to the number of genes shared between a MP-GO term pair, dot color represents the ratio between the number of shared genes over the total number of genes in the GO signature. **B)** Pairwise correlation coefficient between MP signature scores in cancer cells from scRNA-seq data (11 samples). **C)** Pairwise correlation coefficient between MP signature scores in cancer cells from CosMx SMI *in situ* transcripts.

**Figure S3. Overview of tumor samples and Fields-of-View (FOVs) selection for CosMx SMI.** For Patient A & Patient B processed with the Cosmx SMI: (Top panels) Overview immunofluorescence image colored by DAPI (blue), pan-cytokeratin (green), CD68 (yellow) and CD45 (red), and annotated for FOVs selection. (Bottom panels) Overview morphological staining (H&E) from an adjacent slide.

**Figure S4. Detailed H&E and immunofluorescence for individual FOVs in CosMx SMI.** Morphological staining (H&E) and immunofluorescent image of individual FOVs colored by DAPI (blue), pan-cytokeratin (green), CD68 (yellow) and CD45 (red), and annotated for infiltrative, nodular or undetermined morphology (patient A and B).

**Figure S5. Prediction of cell types in CosMx SMI *in situ* transcripts. A)** UMAP embeddings for scRNA-seq samples from Yerly et al. (2022) **B)** Based on the reference expression profiles of the data in A) – plus the expression profile for neutrophils derived from Zilionis et al. (2019) – the inSituType algorithm was applied to transfer labels to the *in situ* transcripts of 4 patients sequenced with CosMx SMI technology. **C)** Normalized average expression (color scale) and percent of expressing cells (dot size) for a panel of marker genes, for each of the cell types defined in the CosMx SMI dataset.

**Figure S6. Morphological complexity of BCCs measured as perimeter/area ratio, in infiltrative vs. nodular Fields-of-View (FOVs).** Morphological complexity measured as perimeter/area ratio on the composite images from Cosmx SMI based on the DAPI (blue) and pan-cytokeratin (green) stainings for FOVs of 2 patients previously annotated as Nodular or Infiltrative. P-value: unpaired t-test.

**Figure S7. Correlation between MPs and nodular-to-infiltrative score (NI score). A)** Density plots for the correlation of MP2 with the NI score (see Methods) in scRNA-seq from 11 BCC patients. **B)** Density plots for the correlation of MP7 with the NI score in scRNA-seq from 11 BCC patients. The correlation coefficient is reported on top of each plot; color scale indicates the density of cells. **C)** Predicted cell types and spatial distribution of MP2 and MP7 in four representative FOVs from Patient B (two nodular, two infiltrative).

**Figure S8. Validation of MP2 and MP7 markers. A)** Expression of *MPPED1* vs. MP2 signature score for cells with nodular, intermediate and infiltrative transcriptomics profiles; and expression of *KRT6A* vs. MP7 signature score for cells with nodular, intermediate and infiltrative transcriptomics profiles. Cells are classified in these three categories by binning the observed NI score range into three equal intervals. **B)** Overview immunofluorescent image stained with DAPI of a human BCC with high intratumoral morphological heterogeneity. White dotted lines indicate tumor areas measured for morphological complexity (perimeter/area) and mean pixel intensity of MP7 (*KRT6A*) and MP2 (*MPPED1*) markers (top left panel). Correlation analysis between MP7 (*KRT6A*) and MP2 (*MPPED1*) markers (bottom left panel). Each dot is colored according to its morphological complexity score (low, medium, high). Illustrative images of low and high complexity tumor areas at higher magnification colored with DAPI (blue), *MPPED1* (green) and *KRT6A* (purple) probes for RNA FISH (right panels). P-value and r: Spearman correlation.

**Figure S9. Meta-program scores in skin cancers of increasing aggressiveness.** Signature scores for MP1-MP7 in scRNA-seq data of nodular and infiltrative BCCs (Yerly et al. 2022); advanced BCCs (Yost et al. 2019); and cSCCs (Ji et al. 2020). Each point represents the average MP score in cancer cells of one sample. cSCC: cutaneous squamous cell carcinoma.

**Figure S10. Morphological features of advanced, therapy-resistant BCCs. A)** Metadata table summarizing advanced BCCs patient’s characteristics. **B)** Morphological complexity measured as perimeter/area ratio in early tumors previously annotated as nodular or infiltrative BCCs by dermatopathologists and in advanced BCCs naive or resistant to HHI treatment. **C)** Representative H&E and immunofluorescent images stained with DAPI (blue) and pan-cytokeratin (grey) of individual advanced BCC samples classified as naive or resistant to treatment. **D)** Patient-matched treatment-naive and treatment-resistant samples shown for representative H&E and immunofluorescent images (left panels), and measured for morphological complexity (measured by perimeter/area ratio; top right panel) and MP7/MP2 ratio (assessed by the ratio of mean pixel intensity of MP7 marker *KRT6A* over mean pixel intensity of MP2 marker *MPPED1;* bottom right panel). P-values: unpaired Student’s *t* test.

**Figure S11. Correlation between MPs and hedgehog (HH) pathway score. A)** Density plots for the correlation of MP2 with the HH pathway score (from KEGG database) in scRNA-seq from 11 BCC patients. **B)** Density plots for the correlation of MP7 with the HH pathway score in scRNA-seq from 11 BCC patients. The correlation coefficient is reported on top of each plot; color scale indicates the density of cells.

**Figure S12. Relation between MP7 and HH pathway markers. A)** Correlation analysis of mean pixel intensity of MP7 marker K*RT6A* and Hedgehog pathway activity G*LI1* at single-cell level across 7 human BCCs. **B)** Representative immunofluorescent images stained with DAPI (blue), pan-cytokeratin (grey), *GLI1* (yellow) and *KRT6A* (purple) probes for RNA FISH. **C)** Correlation analysis of mean pixel intensity of MP7 marker *KRT6A* and Hedgehog pathway activity marker *GLI1* at single-cell level for individual patients. R and p-values: Spearman correlation.

**Figure S13. Morphological complexity in ulcerated BCCs. A)** Representative scan of H&E images of an ulcerated BCC sample in which (A) represents an area “far from ulceration” and (B) an area “close to ulceration”. White dotted lines depict the basement membrane zone (BMZ). **B)** Mean tumor complexity in area far and close to ulceration (N≥6 tumor structures per condition of 21 different BCC samples. Horizontal bars indicate the mean ± SD, p-values were calculated by unpaired two-sided Student’s *t* test. **C)** Representative scan of immunofluorescent images of an ulcerated BCC sample colored with DAPI (blue), *MPPED1* (green) and *KRT6A* (purple) probes for RNA FISH. (A) represents an area “far from ulceration” and (B) an area “close to ulceration. White dotted lines depict the BMZ. D) MP7/MP2 ratio assessed by the ratio of mean pixel intensity of MP7 marker *KRT6A* over mean pixel intensity of MP2 marker *MPPED1.* P-value: unpaired Student’s *t* test.

**Figure S14. Overview of tumor samples and Fields-of-View (FOVs) selection for CosMx SMI.** For Patient C & Patient D processed with the Cosmx SMI: (Top panels) Overview immunofluorescence image colored by DAPI (blue), pan-cytokeratin (green), CD68 (yellow) and CD45 (red), and annotated for FOVs selection. (Bottom panels) Overview morphological staining (H&E) from an adjacent slide.

**Figure S15. Detailed H&E and immunofluorescence for individual FOVs in CosMx SMI.** Morphological staining (H&E) and immunofluorescent image of individual FOVs colored by DAPI (blue), pan-cytokeratin (green), CD68 (yellow) and CD45 (red), and annotated for baseline, wound_far_wound_close and unwound condition (patient C and D). Red arrow heads indicate direction to the wound. N/A: Not available.

**Figure S16. Morphological and transcriptional changes upon BCC wounding. A)** H&E staining for patients 3 and 4 at first biopsy (baseline), at day 7 far from the first biopsy (unwounded), and at day 7 on the same site as the first biopsy (wounded). **B)** Morphological complexity, measured as perimeter/area ratio, for multiple areas of four BCC patients, in the three indicated conditions. **C)** UCell signature score for MP2 and MP7 meta-programs in scRNA-seq from two BCC patients, in the three indicated conditions. **D)** Differentially expressed genes between Wound vs. Baseline/Unwound samples for Patient 2 (data for the other patient in Figure 4). Genes contained in MP2 and MP7 are highlighted blue and red respectively.

**Figure S17. Compositional and transcriptional changes in the TME upon wounding. A)** Compositional changes in the TME upon wounding, measured in a scRNA-seq cohort. For each cell type, the enrichment score is calculated as the log-ratio of the cell type frequency in wounded vs. baseline/unwounded samples (N=2). **B)** Signature scores for the gene sets derived by differential expression analysis in wounded vs. baseline samples, for fibroblasts (CAF), monocyte/macrophages (MoMac), all dendritic cells (panDC) and pericytes, evaluated in 11 nodular and infiltrative BCC scRNA-seq samples. **C)** Wound-responding CAF score in a spatial transcriptomics dataset, averaged by FOVs from baseline, unwounded and wounded samples (far or close to the wound). **D)** Wound-responding CAF score calculated in intratumoral stromal spatial spots from a GeoMx RNA experiment (Yerly et al. 2022), where BCC areas were previously annotated as nodular or infiltrative. p-values: Wilcoxon rank-sum test.

## Methods

### RESOURCE AVAILABILITY

#### Materials availability

Further information and requests for reagents should be directed to François Kuonen. This study did not generate new unique reagents.

#### Data availability

Single-cell RNA sequencing data generated in this study have been deposited in the Gene Expression Omnibus database under accession codes GSE181907 and GSE266665. Spatial transcriptomics data (CosMx SMI) are available from Zenodo under DOI: <u>10.5281/zenodo.14330690</u>.

This paper also analyzes existing, publicly available data. The accession numbers for the datasets are listed in the Resources Table.

#### Code availability

The code to reproduce the computational analyses presented in this manuscript is available at the following repository: https://github.com/CarmonaLab/Yerly_BCC_wounding.

### EXPERIMENTAL MODEL AND STUDY PARTICIPANT DETAILS

#### Clinical samples

Studies were approved by the institutional review board of Lausanne University Hospital CHUV, and the local ethics committee, in accordance with the Helsinki Declaration (CER-VD 2020-02204 and 2021-01984). Written informed consent was obtained from each patient. Based on the principle of voluntary participation, no compensation was provided to the patients in this study. Samples for scRNA-seq were collected from residual material of diagnostic biopsy or surgical piece and processed immediately after resection. Samples for RNA FISH were obtained from formalin-fixed paraffin-embedded tumor tissues. BCC samples referred to as “nodular” and “ulcerated” were identified unbiasedly based on the histopathological report registered between 2022 and 2023, and corresponding H&E slides were collected. Sample sizes are mentioned in the relative results section. Patient characteristics are available in **Table S1**.

### METHOD DETAILS

#### Tissue dissociation and single cell sorting

Fresh tumor tissues were transported in Medium 154CF (Gibco^TM^, M-154CF-500) on ice to preserve viability. The tumor tissues were then minced with scalpels to pieces < 1 mm^3^ and transferred in 10 mL Medium 154CF with collagenase (5 mg/mL, (Gibco^TM^, 17018029)) to digest the tissue. The mixture was incubated for 90 min at 37°C with manual shaking every 10 min. Then, 5 mL 0.05 % trypsin-EDTA (Gibco^TM^, 25300-054) was added to the mixture and the digestion cocktail was incubated again for 10 min at 37°C. After incubation, the mixture was centrifuged at 500 g for 5 min at 4°C. 5 mL 10% fetal bovine serum (FBS, (VWR, S181B-500)) was added to the pellet to inhibit the digestion enzymes. The mixture was centrifuged at 500 g for 5 min at 4°C. Cells were suspended in 10 mL MACS buffer (2% FBS, 2 mM EDTA (Promega, V4231) diluted in phosphate-buffered saline (PBS) (Bichsel, 100 0 324). The cell suspension was then filtered through a 70-µm cell strainer (Plexus-Santé, PL000223). Afterwards, the cells were stained with SYTO^TM^ 13 Green Fluorescent Nucleic Acid Stain (Invitrogen, S7575) and SYTOX^TM^ Orange Nucleic Acid Stain (Invitrogen, S11368) for viability assessment. Stained cells were suspended in MACS buffer. Single living cells (SYTO^TM^ green^Pos^, SYTOX^TM^ orange^Neg^) were then FACS sorted and collected in collecting medium (10% FBS, 2 mM EDTA in PBS). Sorted cells were centrifuged at 500 g for 5 min at 4°C. The cells were then suspended in 10% FBS and counted. The cell suspension was centrifuged at 500 g for 5 min at 4°C. Cells were suspended in the appropriate volume of 10% FBS to obtain a final concentration of approximately 1,000 cells/µl.

#### scRNA-seq processing (Emulsion / Library preparation / Sequencing)

Single-cell mRNA capture and sequencing were performed immediately after reaching the optimal cell suspension by the Lausanne Genomic Technologies Facility (GTF, https://wp.unil.ch/gtf/) using the Chromium Next GEM single cell 3’ v3.1 reagent kit (10x Genomics) following the manufacturer’s protocol. The target number of captured cells was 10,000 per sample. Sequencing libraries were prepared per the manufacturer’s protocol. Sequencing was performed on Illumina NovaSeq 6000 v1.5 (Novaseq Control Software v.1.7.5 for BCC6 and BCC7 or v.1.8.1 for Baseline1/2, Wound1/2 and Unwound1/2) flow cells for 28-10-10-90 cycles (read1 – index i7 – index i5 – read2) with 1% PhiX spike in at a median depth of 23,490 reads/cell for BCC6, 18,397 reads/cell for BCC7, 25,111 reads/cell for Baseline1, 22,592 reads/cell for Wound1, 25,463 reads/cell for Unwound1, 19,049 reads/cell for Basline2, 17,550 reads/cell for Wound2, 16,983 reads/cell for Unwound2. Sequencing data were demultiplexed using the bcl2fastq2 Conversion Software (v. 2.20, Illumina) and primary data analysis performed with 10x Genomics Cell Ranger (v.6.0.0 for BCC6 and BCC7 and v.7.1.0 for Baseline1/2, Wound1/2 and Baseline1/2). The count function allowed us to demultiplex sequencing reads to individual cells, to align the reads to the human GRCh38 genome reference (refdata-gex-GRCh38-2020-A), and to generate filtered matrices of gene counts by cell barcodes.

#### scRNA-seq data processing and quality control

Bioinformatics analyses for scRNA-seq data generated in this study, as well as meta-analyses of previously published datasets (GSE181907^12^, E-MTAB-13085^16^, GSE144236^22^, GSE123814^21^), were performed using the Seurat framework^74^. To make samples from different sources comparable, we harmonized gene names to a common dictionary of Ensembl BioMart gene symbols^75^. Low-quality transcriptomes were excluded from the analysis by filtering out cells with extreme numbers of UMI counts (<600 and >25000), number of detected genes (<500 and >6000), more than 20% of mitochondrial genes, or more than 15% of genes associated with tissue dissociation^76^. Dimensionality reduction was performed using the top 2000 variable genes, and subsequently applying principal component analysis (PCA). The 30 PCA components with highest explained variance were used for unsupervised clustering and for further dimensionality reduction by Uniform Manifold Approximation and Projection (UMAP). Preliminary, automated cell type annotation was obtained using the scGate method and the “TME_HiRes” model^77^, subsequently refining annotations by majority vote of the automated labels in unsupervised clusters (**Fig. S1**). As scGate does not provide models for cancer cells, malignant and normal keratinocytes were identified as outlined in the following section.

#### Identification of malignant cells from scRNA-seq data

We employed a three-tier procedure to distinguish malignant cells from normal keratinocytes, based on *i)* patient mixing by cell type, *ii)* BCC signature scoring and *iii)* copy-number variation (CNV). Patient mixing, quantified by the Local Inverse Simpson’s score (LISI) score^78^, aims at evaluating how well mixed cells from different samples are when embedded together. Because cancer cells tend to have altered gene regulation and abnormal expression, they can be expected to display more sample-specific features and, as a consequence, lower LISI score (see e.g. **Fig S1B** and **S1F**). Second, specific cancer types can display patterns of genes which are consistently differentially expressed compared to healthy cells. In this study, we applied a signature composed of genes previously identified as overexpressed in BCC (*PTCH1*, *GLI1*, *GLI2*, *SPON2*, *HHIP*, *MYCN*) and scored this gene set using UCell^79^ (see e.g. **Fig. S1C** and **S1G**). Third, we applied inferCNV^80^ for copy-number variation analysis to detect chromosomal regions with altered expression patterns compared to reference transcriptomes. Clustering of CNV profiles of cancer cells and normal keratinocytes (using T cells as a reference transcriptome) confirmed that malignant cells had consistent alterations that allowed distinguishing them from healthy keratinocytes (see e.g. **Fig. S1D** and **S1H**).

#### Characterization of cancer gene programs by NMF

To discover gene programs across multiple BCC samples, we applied non-negative matrix factorization (NMF) on 11 samples for which at least 100 cancer cells were available. We implemented our multi-sample NMF discovery algorithm in a R package (GeneNMF), available on the CRAN repository at: https://cran.r-project.org/package=GeneNMF. We describe the algorithm in detail in the “GeneNMF package” methods section further below, but briefly, GeneNMF performs factorization of individual samples and for multiple values of the target number of gene programs. Then, it determines clusters of similar programs which are consistently identified across multiple samples. The consensus signatures of such clusters of programs are termed meta-programs (MP). We ran GeneNMF with k=4:9 (number of target NMF components for each dataset between 4 to 9) on the top 2000 variable genes across samples, and asked for 10 target MPs with min.confidence=0.3. We further filtered GeneNMF results by dropping MPs with fewer than 5 genes, negative average silhouette coefficient, or sample coverage lower than 40%. On the 7 high-confidence MPs, we calculated MP signatures scores by applying the UCell method^79^ and determined differentially expressed genes (DEG) between the top 20% and bottom 20% scoring cells; we only retained genes from the MP sets that were also found among the DEGs, to construct gene sets with specific expression for each MP.

#### Nodular-to-infiltrative (NI) signature for single-cell data

To place cells on a nodular-to-infiltrative axis, we used previously published signatures of nodular and infiltrative cancer cell niches (T^inf^ and T^nod^ respectively) derived from GeoMx spatial transcriptomics^12^. We calculated UCell scores for the T^inf^ and T^nod^ gene sets, and combined them into a single “Nodular-to-infiltrative score” defined as NI.score = UCell(T^inf^) – UCell(T^nod^). This score varies between –1 (fully nodular) to +1 (fully infiltrative).

To identify globally non-invasive and invasive BCC tumors, samples from 11 patients having at least 100 cancer cells were classified using the following criteria. The average NI score by patient was calculated on their cancer cells. Based on the average NI score, patients were split into two groups of equal size: patients with an average NI score above the median were classified as invasive BCC, while those with an average NI score below the median were classified as non-invasive BCC (**Fig. S8A**).

#### Hedgehog pathway signatures for single-cell data

To quantify the activity of the hedgehog (HH) signaling pathway, we obtained the KEGG_HEDGEHOG_SIGNALING_PATHWAY signature from MSigDB^81^. Additionally, we also evaluated a signature of HH targets, consisting of *GLI1*, *HHIP* and *PTCH1*. Per-cell signature scores were obtained by scoring single-cell transcriptomics data using the UCell package^82^.

#### Definition and scoring of “wound-responding CAF” signature

Cancer-associated fibroblasts (CAF) were identified using the scGate ^77^ method with specific fibroblast markers, excluding pericytes (*PDGFA^+^, FBLN1^+^, FBLN2^+^, COL5A1^+^, LUM^+^, CFD^+^, RGS5^-^, NOTCH3^-^, MYOT^-^*) and aggregated into a pseudo-bulk expression matrix for each sample.

Paired differential expression analysis was performed using the DESeq2 package^83^ on pseudo-bulk samples, comparing wounded vs. unwounded and baseline conditions. Genes with a log2 fold change > 0.5 and adjusted p-value < 0.05 were identified as part of the wound signature, while those with a log2 fold change < –0.5 and adjusted p-value < 0.05 constituted the unwound/baseline signature (**Table S3**).

These signatures were scored using UCell^82^ on CAFs from retrospective cohort single-cell samples (GSE181907^12^, E-MTAB-13085^16^, GSE144236^22^, GSE123814^21^), previously classified by scGate^77^. The “wound-responding CAF” score was calculated by subtracting the unwound/baseline signature score from the wound signature score in each cell, defined as: Wound-responding CAF score = UCell(CAF^wound^) – UCell(CAF^unwound/baseline^).

The score was computed on fibroblasts from different datasets: non-cancer, adjacent skin samples (“Normal adjacent”, N=6) obtained from GSE144236, Hedgehog inhibition (HHI)-therapy resistant BCC (N=7, samples from ^21^; GSE123814), squamous cell carcinomas (N=5, samples from^22^; GSE144236), and BCC tumors (N=11, samples from GSE181907^12^ and E-MTAB-13085^16^).

#### Immunofluorescence and Fluorescence *In Situ* Hybridization (FISH) co-detection

FFPE skin blocks were cut and 5-µm sections were mounted onto Superfrost Plus® microscope slides. FFPE skin sections were heated 10 min at 60°C. Slides were deparaffinized in 2 xylene baths of 3 min, then rehydrated in an ethanol gradient from 100% EtOH (2 baths of 3 min), followed by 95% EtOH (3 min) and 70% EtOH (3 min). Slides were then washed in distilled water. RNAscope® Hydrogen Peroxide (ACD, 323100) was added to the tissue sections and slides were incubated for 10 min at room temperature. Slides were then washed for 5 min in distilled water followed by 5 min in PBS 1X. Antigen retrieval was done in 1X Co-detection Target Retrieval buffer (ACD, 323180) pre-warmed at 100°C for 15 min. Slides were then washed twice 2 min in distilled water and 2 min in 1X Phosphate Buffered Saline with 0.1% Tween-20 (PBS-T) (PBS: Bichsel, 100 0 324, Tween-20: Sigma, P1379-500ML). Each tissue section was then covered with the primary antibody solution (Monoclonal antibody to human cytokeratin (pan) (clone Lu-5) (BMA Biomedicals, T-1302) diluted 1:250 in Co-Detection Antibody Diluent (ACD, 323180)) and incubated 2 hours at room temperature in a humidified box. Following primary antibody incubation, slides were washed twice in PBS-T for 2 min. Slides were then fixed for 30 min in Buffered Zinc Formalin (Thermo Scientific, 5701ZF) at room temperature. After fixation, slides were washed four times in PBS-T for 2 min. Protease treatment, probe hybridization, amplifications and stainings were performed with the RNAscope® Multiplex Fluorescent Reagent Kit v2 Assay kit (ACD, 323100) according to the manufacturer’s instructions. After the final *in situ* hybridization horseradish peroxidase (HRP)-blocker step, each tissue section was covered with the secondary antibody dilution (F(ab’)2-Goat anti-Mouse IgG (H+L) Cross-Adsorbed Secondary Antibody, Alexa Fluor™ 488 (Thermo Fisher Scientific, A-11017) diluted 1:500 in Co-Detection Antibody Diluent) for 30 min at room temperature. Slides were then washed twice for 2 min in PBS-T. Finally, slides were mounted with mounting medium with DAPI-Aqueous Fluoroshield (abcam, ab104139). The slides were acquired with Pannoramic 250 slide scanner and processed using SlideViewer 2.5.0.143918 and (Fiji Is Just) ImageJ softwares. The mean pixel intensity of the different FISH stainings was measured in individual cells with imaging processing software package (Fiji Is Just) ImageJ. Chosen areas and distribution over samples are described in the legend of the corresponding figures.

#### Tumor complexity

Tumor complexity was calculated by annotating tumor edges on BCC sample images, and by dividing the perimeter by the area of tumoral structures.

#### CosMx SMI – sample preparation

FFPE skin blocks were cut and 2 consecutive sections of 5-µm were mounted onto individual Superfrost Plus® microscope slides. The tumor sections from four different patients were arranged on 2 slides (Patient A and C on slide 1, Patient B and D on slide 2). Hematoxylin and eosin staining was performed on the first section; the second, adjacent section was prepared for CosMx SMI as follows. To ensure FFPE tissue adherence to the glass slides, they were baked overnight at 60°C. Samples underwent deparaffinization, proteinase K digestion, and heat-induced epitope retrieval (HIER) procedures to expose target RNAs and epitopes using Leica Bond Rx system. Proteinase K (3 μg/ml; ThermoFisher) incubation at 40°C for 30 min and HIER at 100°C for 15 min in Leica buffer ER1 conditions were used. The samples were rinsed with diethyl pyrocarbonate (DEPC)-treated water (DEPC H2O) twice before incubating with 1:1000 diluted fiducials (Bangs Laboratory) in 2X SSCT (2X saline sodium citrate, 0.001% Tween 20) solution for 5 min at room temperature. Excess fiducials were removed by rinsing the samples with 1X phosphate buffered saline (PBS), followed by fixation with 10% neutral buffered formalin (NBF) for 5 min at room temperature. Fixed samples were rinsed with Tris-glycine buffer (0.1M glycine, 0.1M Tris-base in DEPC H2O) and 1X PBS for 5 min each before blocked using 100 mM N-succinimidyl acetate (NHS-acetate, ThermoFisher) in NHS-acetate buffer (0.1M NaP, 0.1% Tween pH 8 in DEPC H2O) for 15 min at room temperature. Prepared samples were rinsed with 2X saline sodium citrate (SSC) for 5 min and then Adhesive SecureSeal Hybridization Chamber (Grace Bio-Labs) was placed to cover the samples. RNA ISH probes (980plex) were denatured at 95°C for 2 min and then placed on ice before preparing the ISH probe mix (1 nM ISH probes, 1X Buffer R, 0.1 U/μL SUPERaseIn™ in DEPC H2O). The ISH probe mix was pipetted into the hybridization chamber and the chamber was sealed using adhesive tape. Hybridization was performed at 37°C overnight to prevent evaporation. After the overnight hybridization, samples were washed twice with 50% formamide (VWR) in 2X SSC at 37°C for 25 min, rinsed twice with 2X SSC for 2 min at room temperature, and then blocked with 100 mM NHS-acetate for 15 min. After blocking, the samples were washed twice using 2X SSC for 2 min at room temperature. Custom-made slide covers were attached to the sample slide to form a flow cell.

#### CosMx SMI – spatial transcriptomics measurement

Target RNA readout on SMI instrument was performed following published protocols^13^. In brief, the assembled flow cell was loaded onto the SMI instrument and the samples were washed with reporter wash buffer to rinse the samples and remove air bubbles. Once the flow cell was loaded onto the instrument, the entire flow cell was scanned and 45 Fields-of-View (FOVs) of size 0.51mm by 0.51mm were placed on each slide for RNA readout. The first RNA readout cycle was initiated by flowing 100 µL of Reporter Pool 1 into the flow cell and incubating for 15 min. After incubation, 1 mL of Reporter Wash Buffer was flowed across the flow cell to wash out the unbound reporter probes, followed by replacing Reporter Wash buffer with Imaging buffer prior to imaging. Nine Z-stack images (0.8 µm step size) of each FOV were acquired and then fluorophores on the reporter probes were UV cleaved and washed off with Strip Wash buffer. This fluidic and imaging procedure was then repeated for the remaining 15 reporter pools, and the 16 cycles of reporter hybridization-imaging was repeated 8 times to increase RNA detection sensitivity. After 9 complete cycles of RNA readout, samples were incubated with a fluorophore-conjugated antibody cocktail against CD298/B2M, PanCK, CD45, and CD3 proteins and DAPI in the same instrument for 1 hour. Nine Z-stack images for 5 channels (4 antibodies and DAPI) were captured after unbound antibodies and DAPI were washed with Reporter washing buffer and flow cell was filled with Imaging buffer.

#### CosMx SMI – image processing and data acquisition

Raw image processing and feature extraction were performed using a Nanostring in-house SMI data processing pipeline^13^ which includes registration, feature detection, and localization. 3D rigid image registration was made using fiducials embedded in the samples matched with the fixed image reference established at the beginning of SMI run to correct for any shift. Secondly, RNA image analysis algorithm was used to identify reporter signature locations in X, Y, and Z axes along with the assigned confidence. The reporter signature locations and the associated features were collated into a single list. Lastly, the XYZ location information of individual target transcript was extracted and recorded in a table by a secondary analysis algorithm, as described in^13^. The Z-stack images of immunostaining and DAPI were used for drawing cell boundaries on the samples. A cell segmentation pipeline using machine learning algorithm^84,85^ was used to accurately assign transcripts to cell locations and subcellular compartments. The transcript profile of individual cells was generated by combining target transcript location and cell segmentation boundaries. The pre-processed spatial transcriptomics data were exported to .rds files, and subsequent analyses were performed in R and Seurat^86^.

#### CosMx SMI – cell type prediction

To predict cell type identities from CosMx *in situ* transcripts, we applied the InSituType algorithm^19^ (see **Fig. S5**). InSituType annotation of a query dataset is based on label transfer from a reference set of annotated transcriptomes. We used a set of 7 confidently-annotated scRNA-seq samples (BCC1 to BCC7, Yerly et al. cohort) as a reference for label transfer, to which we manually added an average expression profile for neutrophils derived from Zilionis et al.^87^. Cells with < 100 detected transcripts and with >2% transcripts coming from negative control probes were labeled as “Low quality” cells.

#### Definition of homotypic and heterotypic cell-cell interactions in spatial transcriptomics data

Homotypic cell-cell interactions consist of pairs of cells of the same type; heterotypic interactions include cells of different cell types. To define cancer cells that participate in homotypic or heterotypic interactions in our CosMx spatial datasets, we first calculated nearest neighbor graphs based on spatial coordinates using the BiocNeighbors package (DOI: 10.18129/B9.bioc.BiocNeighbors). We limited the neighbor search between cell centroids to a maximum distance D, corresponding to 2.5 times the average cell diameter in the CosMx dataset. If ≥90% of the cells within distance D of a given cancer cells were also cancer cells, the cancer cell was labeled as “homotypic”; otherwise it was labeled as “heterotypic”.

#### Graph-based analysis of cell-cell interactions

To summarize and visualize the interactome between cell types in spatial transcriptomics data, we applied a graph-based approach inspired by previous work^88^. For any given cell, we calculated the physical euclidean distance to the nearest neighbor of each cell type; averaging by cell type and by experimental condition allowed us to compile distance matrices of mean and standard deviation for all pairs of cell types. Based on these matrices, we applied *igraph*^89^ with the Kamada-Kawai layout algorithm to construct graph representations of pairwise cell type interactions. In this representation, edge lengths approximate the mean cell type distances (constrained by a model of springs on a plane); edge widths are proportional to the standard deviation of the distances; and node sizes are proportional to the abundance of each cell type in a given condition.

#### Signature scoring in spatial data and correlation with wound proximity

Gene signature scoring (for MPs, CAF and HH signatures) was performed on CosMx data by applying the UCell algorithm with default parameters on the *in situ* transcripts. Genes not included in the CosMx 6k panel were excluded from all signatures. A wound axis was established for each FOV by assigning an approximate directionality of the wound site (by steps of 45 degrees); a “wound proximity” value was then calculated for each cell based on its location on the wound axis within each FOV. Correlations between signature scores and wound proximity were calculated in terms of Spearman correlation coefficient.

#### GeneNMF package

We developed an R package, GeneNMF, to streamline the identification of gene programs from scRNA-seq data using non-negative matrix factorization (NMF). NMF is a dimensionality reduction technique that aims at decomposing a high-dimensional data matrix **A** into the product of two matrices **W** and **H** such that **A ≅ WH**, with the constraints that all elements in the three matrices should be zero or positive. In the case of scRNA-seq data, **A** is the observed gene expression matrix of size g x c (where g is the number of genes and c the number of cells); **H** is the embedding matrix of size k x c (where k is the number of gene programs) containing the coordinates of cells in the low dimensional space; and **W** is the feature loading matrix of size g x k, containing the coefficients of each gene in each program.

Given a list of expression matrices, one for each sample, GeneNMF performs NMF decomposition individually for each sample using the RcppML library (https://CRAN.R-project.org/package=RcppML)^90^. Critical parameters are the set of genes used as input (by default the top 2000 variable genes); the minimum and maximum average log-expression value for including genes in the decomposition; and the number of components **k**, corresponding to the target number of gene programs for a single run. The multiNMF() function of GeneNMF allows specifying a range of values for **k**, prompting the method to generate a NMF decomposition for each value of **k**, for each sample. The method then extracts the most program-specific genes using the extractFeatures() methods implemented in the “NMF” package^91^. Similarly to the approach by Gavish et *al.*^17^, GeneNMF then clusters gene programs by the fraction of shared genes, as can be quantified using the Jaccard similarity coefficient. The consensus of the gene sets in any given cluster of programs is termed meta-program (MP). Several metrics are generated by GeneNMF for each MP. In particular, the average Jaccard index informs on the internal consistency of individual programs within a MP; the silhouette width coefficient quantifies whether a MPs is specific, or whether it shares genes with other MPs; sample coverage quantifies the proportion of samples in which the MP was detected. These metrics can be used downstream to evaluate the quality of the decomposition, assess properties of individual MPs, and filter out inconsistent or redundant MPs.

We note that NMF has been previously applied for the decomposition of single-cell data^17,92,93^. In particular, the consensus NMF (cNMF) method^93^ addressed a potential limitation of NMF, namely its reliance on stochastic optimization algorithms and consequent non-deterministic nature. cNMF introduced the idea of performing multiple decomposition replicates from different initialization seeds, –and then calculating a consensus of the resulting multiple solutions. Inspired by the work of Gavish et *al.*^17^, GeneNMF extends the consensus NMF approach to span multiple values of **k,** a crucial parameter that determines the target number of programs for each NMF run, and which is generally a priori unknown. Importantly, GeneNMF can interact directly with the Seurat infrastructure, making it very straightforward to incorporate in analysis pipelines for single-cell analysis.

---

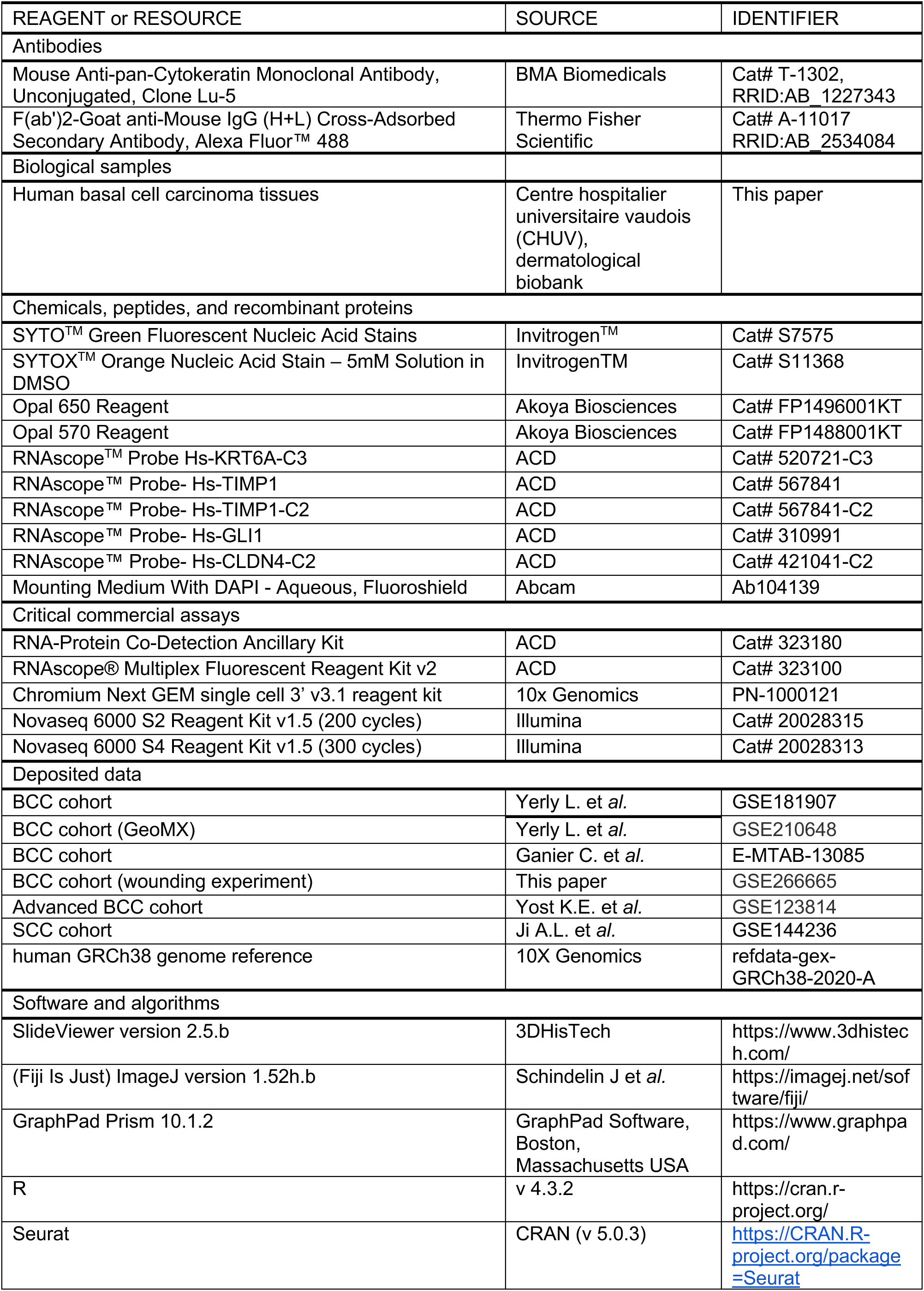
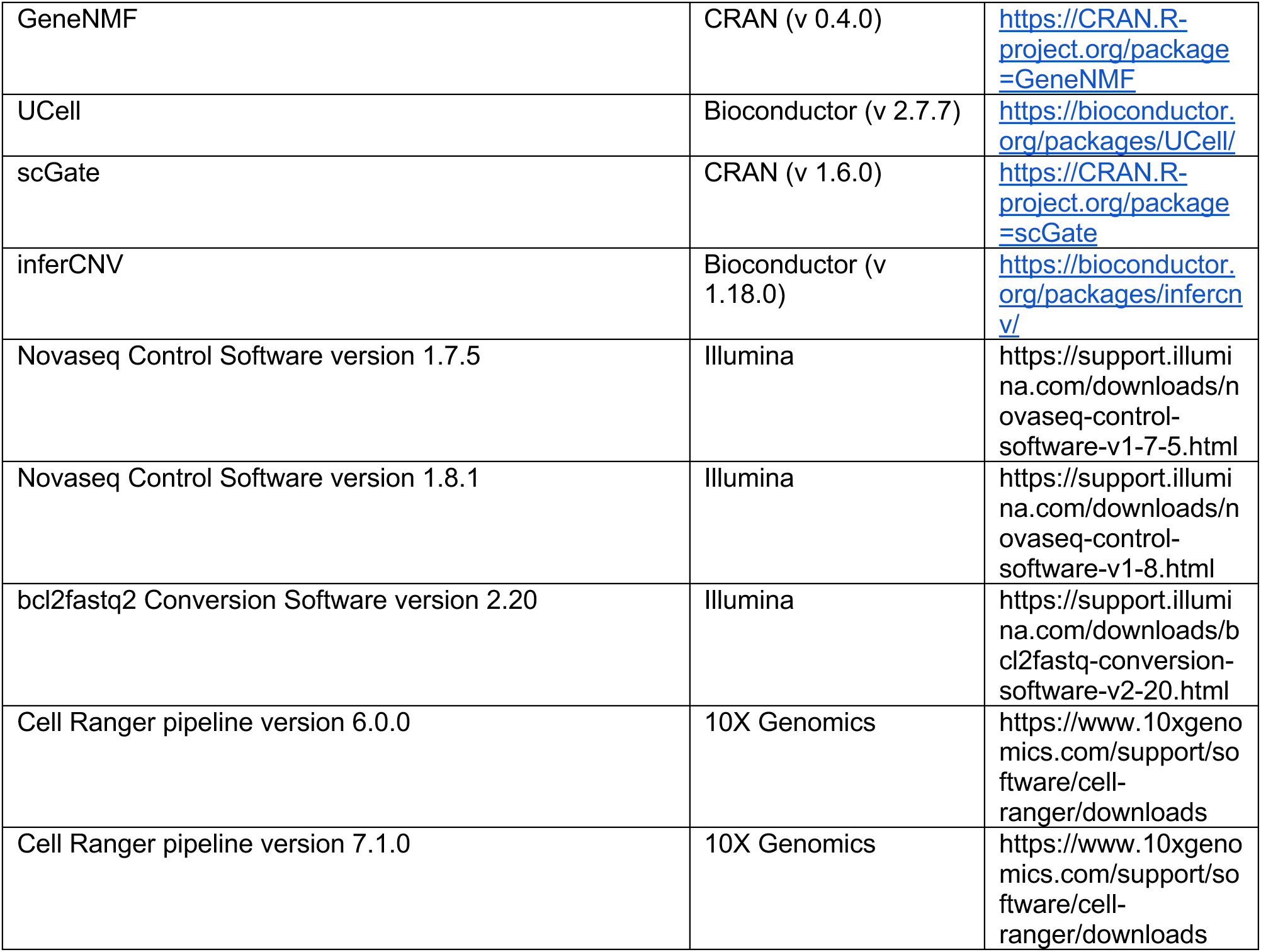
RESOURCES TABLE.

RESOURCES TABLE
| REAGENT or RESOURCE | SOURCE | IDENTIFIER |
| --- | --- | --- |
| **Antibodies** |  |  |
| Mouse Anti-pan-Cytokeratin Monoclonal Antibody, Unconjugated, Clone Lu-5 | BMA Biomedicals | Cat# T-1302, RRID:AB_1227343 |
| F(ab')<sub>2</sub>-Goat anti-Mouse IgG (H+L) Cross-Adsorbed Secondary Antibody, Alexa Fluor™ 488 | Thermo Fisher Scientific | Cat# A-11017 RRID:AB_2534084 |
| **Biological samples** |  |  |
| Human basal cell carcinoma tissues | Centre hospitalier universitaire vaudois (CHUV), dermatological biobank | This paper |
| **Chemicals, peptides, and recombinant proteins** |  |  |
| SYTO™ Green Fluorescent Nucleic Acid Stains | Invitrogen™ | Cat# S7575 |
| SYTOX™ Orange Nucleic Acid Stain – 5mM Solution in DMSO | Invitrogen™ | Cat# S11368 |
| Opal 650 Reagent | Akoya Biosciences | Cat# FP1496001KT |
| Opal 570 Reagent | Akoya Biosciences | Cat# FP1488001KT |
| RNAscope™ Probe Hs-KRT6A-C3 | ACD | Cat# 520721-C3 |
| RNAscope™ Probe- Hs-TIMP1 | ACD | Cat# 567841 |
| RNAscope™ Probe- Hs-TIMP1-C2 | ACD | Cat# 567841-C2 |
| RNAscope™ Probe- Hs-GLI1 | ACD | Cat# 310991 |
| RNAscope™ Probe- Hs-CLDN4-C2 | ACD | Cat# 421041-C2 |
| Mounting Medium With DAPI - Aqueous, Fluoroshield | Abcam | Ab104139 |
| **Critical commercial assays** |  |  |
| RNA-Protein Co-Detection Ancillary Kit | ACD | Cat# 323180 |
| RNAscope® Multiplex Fluorescent Reagent Kit v2 | ACD | Cat# 323100 |
| Chromium Next GEM single cell 3' v3.1 reagent kit | 10x Genomics | PN-1000121 |
| Novaseq 6000 S2 Reagent Kit v1.5 (200 cycles) | Illumina | Cat# 20028315 |
| Novaseq 6000 S4 Reagent Kit v1.5 (300 cycles) | Illumina | Cat# 20028313 |
| **Deposited data** |  |  |
| BCC cohort | Yerly L. et al. | GSE181907 |
| BCC cohort (GeoMX) | Yerly L. et al. | GSE210648 |
| BCC cohort | Ganier C. et al. | E-MTAB-13085 |
| BCC cohort (wounding experiment) | This paper | GSE266665 |
| Advanced BCC cohort | Yost K.E. et al. | GSE123814 |
| SCC cohort | Ji A.L. et al. | GSE144236 |
| human GRCh38 genome reference | 10X Genomics | refdata-gex-GRCh38-2020-A |
| **Software and algorithms** |  |  |
| SlideViewer version 2.5.b | 3DHisTech | https://www.3dhistech.com/ |
| (Fiji Is Just) ImageJ version 1.52h.b | Schindelin J et al. | https://imagej.net/software/fiji/ |
| GraphPad Prism 10.1.2 | GraphPad Software, Boston, Massachusetts USA | https://www.graphpad.com/ |
| R | v 4.3.2 | https://cran.r-project.org/ |
| Seurat | CRAN (v 5.0.3) | https://CRAN.R-project.org/package=Seurat |
| GeneNMF | CRAN (v 0.4.0) | https://CRAN.R-project.org/package=GeneNMF |
| UCell | Bioconductor (v 2.7.7) | https://bioconductor.org/packages/UCell/ |
| scGate | CRAN (v 1.6.0) | https://CRAN.R-project.org/package=scGate |
| inferCNV | Bioconductor (v 1.18.0) | https://bioconductor.org/packages/infercnv/ |
| Novaseq Control Software version 1.7.5 | Illumina | https://support.illumina.com/downloads/novaseq-control-software-v1-7-5.html |
| Novaseq Control Software version 1.8.1 | Illumina | https://support.illumina.com/downloads/novaseq-control-software-v1-8.html |
| bcl2fastq2 Conversion Software version 2.20 | Illumina | https://support.illumina.com/downloads/bcl2fastq-conversion-software-v2-20.html |
| Cell Ranger pipeline version 6.0.0 | 10X Genomics | https://www.10xgenomics.com/support/software/cell-ranger/downloads |
| Cell Ranger pipeline version 7.1.0 | 10X Genomics | https://www.10xgenomics.com/support/software/cell-ranger/downloads |

## References

1. Cameron, M. C. et al. Basal cell carcinoma. J. Am. Acad. Dermatol. 80, 303–317 (2019).

2. Crowson, A. N. Basal cell carcinoma: biology, morphology and clinical implications. Mod. Pathol. 19, S127–S147 (2006).

3. Armstrong, L. T. D., Magnusson, M. R. & Guppy, M. P. B. Risk factors for recurrence of facial basal cell carcinoma after surgical excision: A follow-up analysis. J. Plast. Reconstr. Aesthet. Surg. 70, 1738–1745 (2017).

4. Piccinno, R., Benardon, S., Gaiani, F. M., Rozza, M. & Caccialanza, M. Dermatologic radiotherapy in the treatment of extensive basal cell carcinomas: a retrospective study. J. Dermatol. Treat. 28, 426–430 (2017).

5. Caccialanza, M., Piccinno, R., Çuka, E., Alberti Violetti, S. & Rozza, M. Radiotherapy of morphea-type basal cell carcinoma: results in 127 cases. J. Eur. Acad. Dermatol. Venereol. 28, 1751–1755 (2014).

6. Epstein, E. H. Basal cell carcinomas: attack of the hedgehog. Nat. Rev. Cancer 8, 743–754 (2008).

7. Sekulic, A. et al. Efficacy and Safety of Vismodegib in Advanced Basal-Cell Carcinoma. N. Engl. J. Med. 366, 2171–2179 (2012).

8. Chang, A. L. S. & Oro, A. E. Initial Assessment of Tumor Regrowth After Vismodegib in Advanced Basal Cell Carcinoma. Arch. Dermatol. 148, 1324 (2012).

9. Bonilla, X. et al. Genomic analysis identifies new drivers and progression pathways in skin basal cell carcinoma. Nat. Genet. 48, 398–406 (2016).

10. Villani, R. et al. Subtype-Specific Analyses Reveal Infiltrative Basal Cell Carcinomas Are Highly Interactive with their Environment. J. Invest. Dermatol. 141, 2380–2390 (2021).

11. Kuonen, F. et al. TGFβ, Fibronectin and Integrin α5β1 Promote Invasion in Basal Cell Carcinoma. J. Invest. Dermatol. 138, 2432–2442 (2018).

12. Yerly, L. et al. Integrated multi-omics reveals cellular and molecular interactions governing the invasive niche of basal cell carcinoma. Nat. Commun. 13, 4897 (2022).

13. He, S. et al. High-plex imaging of RNA and proteins at subcellular resolution in fixed tissue by spatial molecular imaging. Nat. Biotechnol. 40, 1794–1806 (2022).

14. Kim, Y. et al. Seq-Scope: repurposing Illumina sequencing flow cells for high-resolution spatial transcriptomics. Nat. Protoc. (2024) doi:10.1038/s41596-024-01065-0.

15. Tirosh, I. & Suva, M. L. Cancer cell states: Lessons from ten years of single-cell RNA-sequencing of human tumors. Cancer Cell 42, 1497–1506 (2024).

16. Ganier, C. et al. Multiscale spatial mapping of cell populations across anatomical sites in healthy human skin and basal cell carcinoma. Proc. Natl. Acad. Sci. 121, e2313326120 (2024).

17. Gavish, A. et al. Hallmarks of transcriptional intratumour heterogeneity across a thousand tumours. Nature 618, 598–606 (2023).

18. He, S. et al. High-plex Multiomic Analysis in FFPE at Subcellular Level by Spatial Molecular Imaging. Preprint at 10.1101/2021.11.03.467020 (2021).

19. Danaher, P. et al. Insitutype: likelihood-based cell typing for single cell spatial transcriptomics. Preprint at 10.1101/2022.10.19.512902 (2022).

20. Hanahan, D. Hallmarks of Cancer: New Dimensions. Cancer Discov. 12, 31–46 (2022).

21. Yost, K. E. et al. Clonal replacement of tumor-specific T cells following PD-1 blockade. Nat. Med. 25, 1251–1259 (2019).

22. Ji, A. L. et al. Multimodal Analysis of Composition and Spatial Architecture in Human Squamous Cell Carcinoma. Cell 182, 497–514.e22 (2020).

23. Winge, M. C. G. et al. Advances in cutaneous squamous cell carcinoma. Nat. Rev. Cancer 23, 430–449 (2023).

24. Boumahdi, S. & De Sauvage, F. J. The great escape: tumour cell plasticity in resistance to targeted therapy. Nat. Rev. Drug Discov. 19, 39–56 (2020).

25. Biehs, B. et al. A cell identity switch allows residual BCC to survive Hedgehog pathway inhibition. Nature 562, 429–433 (2018).

26. Sánchez-Danés, A. et al. A slow-cycling LGR5 tumour population mediates basal cell carcinoma relapse after therapy. Nature 562, 434–438 (2018).

27. Kuonen, F. et al. Loss of Primary Cilia Drives Switching from Hedgehog to Ras/MAPK Pathway in Resistant Basal Cell Carcinoma. J. Invest. Dermatol. 139, 1439–1448 (2019).

28. Kuonen, F. et al. c-FOS drives reversible basal to squamous cell carcinoma transition. Cell Rep. 37, 109774 (2021).

29. Joost, S. et al. Single-Cell Transcriptomics Reveals that Differentiation and Spatial Signatures Shape Epidermal and Hair Follicle Heterogeneity. Cell Syst. 3, 221–237.e9 (2016).

30. Peris, K. et al. European consensus-based interdisciplinary guideline for diagnosis and treatment of basal cell carcinoma—update 2023. Eur. J. Cancer 192, 113254 (2023).

31. Altamura, D. et al. Dermatoscopy of basal cell carcinoma: Morphologic variability of global and local features and accuracy of diagnosis. J. Am. Acad. Dermatol. 62, 67–75 (2010).

32. Wietecha, M. S. et al. Phase-specific signatures of wound fibroblasts and matrix patterns define cancer-associated fibroblast subtypes. Matrix Biol. 119, 19–56 (2023).

33. MacCarthy-Morrogh, L. & Martin, P. The hallmarks of cancer are also the hallmarks of wound healing. Sci. Signal. 13, eaay8690 (2020).

34. Fine, J.-D., Johnson, L. B., Weiner, M., Li, K.-P. & Suchindran, C. Epidermolysis bullosa and the risk of life-threatening cancers: The National EB Registry experience, 1986-2006. J. Am. Acad. Dermatol. 60, 203–211 (2009).

35. Özyazgan, İ. & Kontaş, O. Previous injuries or scars as risk factors for the development of basal cell carcinoma. Scand. J. Plast. Reconstr. Surg. Hand Surg. 38, 11–15 (2004).

36. Hartnett, L. & Egan, L. J. Inflammation, DNA methylation and colitis-associated cancer. Carcinogenesis 33, 723–731 (2012).

37. Valenzuela, M. A., Canales, J., Corvalán, A. H. & Quest, A. F. G. Helicobacter pylori-induced inflammation and epigenetic changes during gastric carcinogenesis. World J. Gastroenterol. 21, 12742–12756 (2015).

38. Antonio, N. et al. The wound inflammatory response exacerbates growth of pre-neoplastic cells and progression to cancer. EMBO J. 34, 2219–2236 (2015).

39. Katikaneni, A. et al. Lipid peroxidation regulates long-range wound detection through 5-lipoxygenase in zebrafish. Nat. Cell Biol. 22, 1049–1055 (2020).

40. Uderhardt, S., Martins, A. J., Tsang, J. S., Lämmermann, T. & Germain, R. N. Resident Macrophages Cloak Tissue Microlesions to Prevent Neutrophil-Driven Inflammatory Damage. Cell 177, 541–555.e17 (2019).

41. Razzell, W., Evans, I. R., Martin, P. & Wood, W. Calcium Flashes Orchestrate the Wound Inflammatory Response through DUOX Activation and Hydrogen Peroxide Release. Curr. Biol. 23, 424–429 (2013).

42. Li, N. Y. et al. Basal-to-inflammatory transition and tumor resistance via crosstalk with a pro-inflammatory stromal niche. Nat. Commun. 15, 8134 (2024).

43. Starace, M. et al. Management of malignant cutaneous wounds in oncologic patients. Support. Care Cancer 30, 7615–7623 (2022).

44. Liu, S. et al. A tissue injury sensing and repair pathway distinct from host pathogen defense. Cell 186, 2127–2143.e22 (2023).

45. Peña, O. A. & Martin, P. Cellular and molecular mechanisms of skin wound healing. Nat. Rev. Mol. Cell Biol. 25, 599–616 (2024).

46. Werner, S., Krieg, T. & Smola, H. Keratinocyte–Fibroblast Interactions in Wound Healing. J. Invest. Dermatol. 127, 998–1008 (2007).

47. Park, S. et al. Tissue-scale coordination of cellular behaviour promotes epidermal wound repair in live mice. Nat. Cell Biol. 19, 155–163 (2017).

48. Gedeon Matoltsy, A. & Viziam, C. B. Further Observations on Epithelialization of Small Wounds. J. Invest. Dermatol. 55, 20–25 (1970).

49. Garlick, J. A. & Taichman, L. B. Fate of human keratinocytes during reepithelialization in an organotypic culture model. Lab. Investig. J. Tech. Methods Pathol. 70, 916–924 (1994).

50. Aragona, M. et al. Defining stem cell dynamics and migration during wound healing in mouse skin epidermis. Nat. Commun. 8, 14684 (2017).

51. Slack, R. J., Macdonald, S. J. F., Roper, J. A., Jenkins, R. G. & Hatley, R. J. D. Emerging therapeutic opportunities for integrin inhibitors. Nat. Rev. Drug Discov. 21, 60–78 (2022).

52. Koivisto, L., Häkkinen, L. & Larjava, H. Re-epithelialization of wounds. Endod. Top. 24, 59–93 (2011).

53. Marsh, D. et al. αvβ6 Integrin Promotes the Invasion of Morphoeic Basal Cell Carcinoma through Stromal Modulation. Cancer Res. 68, 3295–3303 (2008).

54. Wietecha, M. S. et al. Activin-mediated alterations of the fibroblast transcriptome and matrisome control the biomechanical properties of skin wounds. Nat. Commun. 11, 2604 (2020).

55. Cangkrama, M., et al. A paracrine activin A–mDia2 axis promotes squamous carcinogenesis via fibroblast reprogramming. EMBO Mol. Med. 12, e11466 (2020).

56. Martin, P. Wound Healing--Aiming for Perfect Skin Regeneration. Science 276, 75–81 (1997).

57. Bansaccal, N. et al. The extracellular matrix dictates regional competence for tumour initiation. Nature 623, 828–835 (2023).

58. Lim, H. et al. Discrepancy between endoscopic forceps biopsy and endoscopic resection in gastric epithelial neoplasia. Surg. Endosc. 28, 1256–1262 (2014).

59. Kim, Y. et al. Histologic diagnosis based on forceps biopsy is not adequate for determining endoscopic treatment of gastric adenomatous lesions. Endoscopy 42, 620–626 (2010).

60. Izikson, L., Seyler, M. & Zeitouni, N. C. Prevalence of Underdiagnosed Aggressive Non-Melanoma Skin Cancers Treated with Mohs Micrographic Surgery: Analysis of 513 Cases. Dermatol. Surg. 36, 1769–1772 (2010).

61. Haws, A. L., Rojano, R., Tahan, S. R. & Phung, T. L. Accuracy of biopsy sampling for subtyping basal cell carcinoma. J. Am. Acad. Dermatol. 66, 106–111 (2012).

62. Mosterd, K. et al. Correlation between histologic findings on punch biopsy specimens and subsequent excision specimens in recurrent basal cell carcinoma. J. Am. Acad. Dermatol. 64, 323–327 (2011).

63. Dik, E. A. et al. Poor Correlation of Histologic Parameters Between Biopsy and Resection Specimen in Early Stage Oral Squamous Cell Carcinoma. Am. J. Clin. Pathol. 144, 659–666 (2015).

64. Daveau, C. et al. Histological grade concordance between diagnostic core biopsy and corresponding surgical specimen in HR-positive/HER2-negative breast carcinoma. Br. J. Cancer 110, 2195–2200 (2014).

65. Fiechter, S. et al. Facial Basal Cell Carcinomas Recurring after Photodynamic Therapy: A Retrospective Analysis of Histological Subtypes. Dermatology 224, 346–351 (2012).

66. Boulinguez, S. et al. Histological evolution of recurrent basal cell carcinoma and therapeutic implications for incompletely excised lesions. Br. J. Dermatol. 151, 623–626 (2004).

67. Menn, H. The Recurrent Basal Cell Epithelioma: A Study of 100 Cases of Recurrent, Re-treated Basal Cell Epitheliomas. Arch. Dermatol. 103, 628 (1971).

68. Lotti, F. et al. Chemotherapy activates cancer-associated fibroblasts to maintain colorectal cancer-initiating cells by IL-17A. J. Exp. Med. 210, 2851–2872 (2013).

69. Wynn, T. Cellular and molecular mechanisms of fibrosis. J. Pathol. 214, 199–210 (2008).

70. Verset, L. et al. Impact of neoadjuvant therapy on cancer-associated fibroblasts in rectal cancer. Radiother. Oncol. 116, 449–454 (2015).

71. Barker, H. E., Paget, J. T. E., Khan, A. A. & Harrington, K. J. The tumour microenvironment after radiotherapy: mechanisms of resistance and recurrence. Nat. Rev. Cancer 15, 409–425 (2015).

72. Kuonen, F., Secondini, C. & Rüegg, C. Molecular Pathways: Emerging Pathways Mediating Growth, Invasion, and Metastasis of Tumors Progressing in an Irradiated Microenvironment. Clin. Cancer Res. 18, 5196–5202 (2012).

73. Watson, S. S. et al. Fibrotic response to anti-CSF-1R therapy potentiates glioblastoma recurrence. Cancer Cell 42, 1507–1527.e11 (2024).

74. Hao, Y. et al. Integrated analysis of multimodal single-cell data. Cell 184, 3573–3587.e29 (2021).

75. Martin, F. J. et al. Ensembl 2023. Nucleic Acids Res. 51, D933–D941 (2023).

76. Van Den Brink, S. C., et al. Single-cell sequencing reveals dissociation-induced gene expression in tissue subpopulations. Nat. Methods 14, 935–936 (2017).

77. Andreatta, M., Berenstein, A. J. & Carmona, S. J. scGate: marker-based purification of cell types from heterogeneous single-cell RNA-seq datasets. Bioinformatics 38, 2642–2644 (2022).

78. Korsunsky, I. et al. Fast, sensitive and accurate integration of single-cell data with Harmony. Nat. Methods 16, 1289–1296 (2019).

79. Andreatta, M. & Carmona, S. J. UCell: Robust and scalable single-cell gene signature scoring. Comput. Struct. Biotechnol. J. 19, 3796–3798 (2021).

80. Tirosh, I. et al. Dissecting the multicellular ecosystem of metastatic melanoma by single-cell RNA-seq. Science 352, 189–196 (2016).

81. Liberzon, A. et al. The Molecular Signatures Database Hallmark Gene Set Collection. Cell Syst. 1, 417–425 (2015).

82. Andreatta, M. & Carmona, S. J. UCell: Robust and scalable single-cell gene signature scoring. Comput. Struct. Biotechnol. J. 19, 3796–3798 (2021).

83. Love, M. I., Huber, W. & Anders, S. Moderated estimation of fold change and dispersion for RNA-seq data with DESeq2. Genome Biol. 15, 550 (2014).

84. Stringer, C., Wang, T., Michaelos, M. & Pachitariu, M. Cellpose: a generalist algorithm for cellular segmentation. Nat. Methods 18, 100–106 (2021).

85. Pachitariu, M. & Stringer, C. Cellpose 2.0: how to train your own model. Nat. Methods 19, 1634– 1641 (2022).

86. Hao, Y. et al. Dictionary learning for integrative, multimodal and scalable single-cell analysis. Nat. Biotechnol. 42, 293–304 (2024).

87. Zilionis, R. et al. Single-Cell Transcriptomics of Human and Mouse Lung Cancers Reveals Conserved Myeloid Populations across Individuals and Species. Immunity 50, 1317–1334.e10 (2019).

88. Watson, S. S. et al. Microenvironmental reorganization in brain tumors following radiotherapy and recurrence revealed by hyperplexed immunofluorescence imaging. Nat. Commun. 15, 3226 (2024).

89. Csárdi, G., et al. igraph for R: R interface of the igraph library for graph theory and network analysis. Zenodo 10.5281/ZENODO.7682609 (2024).

90. DeBruine, Z. J., Pospisilik, J. A. & Triche, T. J. Fast and interpretable non-negative matrix factorization for atlas-scale single cell data. Preprint at 10.1101/2021.09.01.458620 (2021).

91. Gaujoux, R. & Seoighe, C. A flexible R package for nonnegative matrix factorization. BMC Bioinformatics 11, 367 (2010).

92. Barkley, D. et al. Cancer cell states recur across tumor types and form specific interactions with the tumor microenvironment. Nat. Genet. 54, 1192–1201 (2022).

93. Kotliar, D. et al. Identifying gene expression programs of cell-type identity and cellular activity with single-cell RNA-Seq. eLife 8, e43803 (2019).

