## Supplementary Figures for "Wounding triggers invasive progression in human basal cell carcinoma"

---

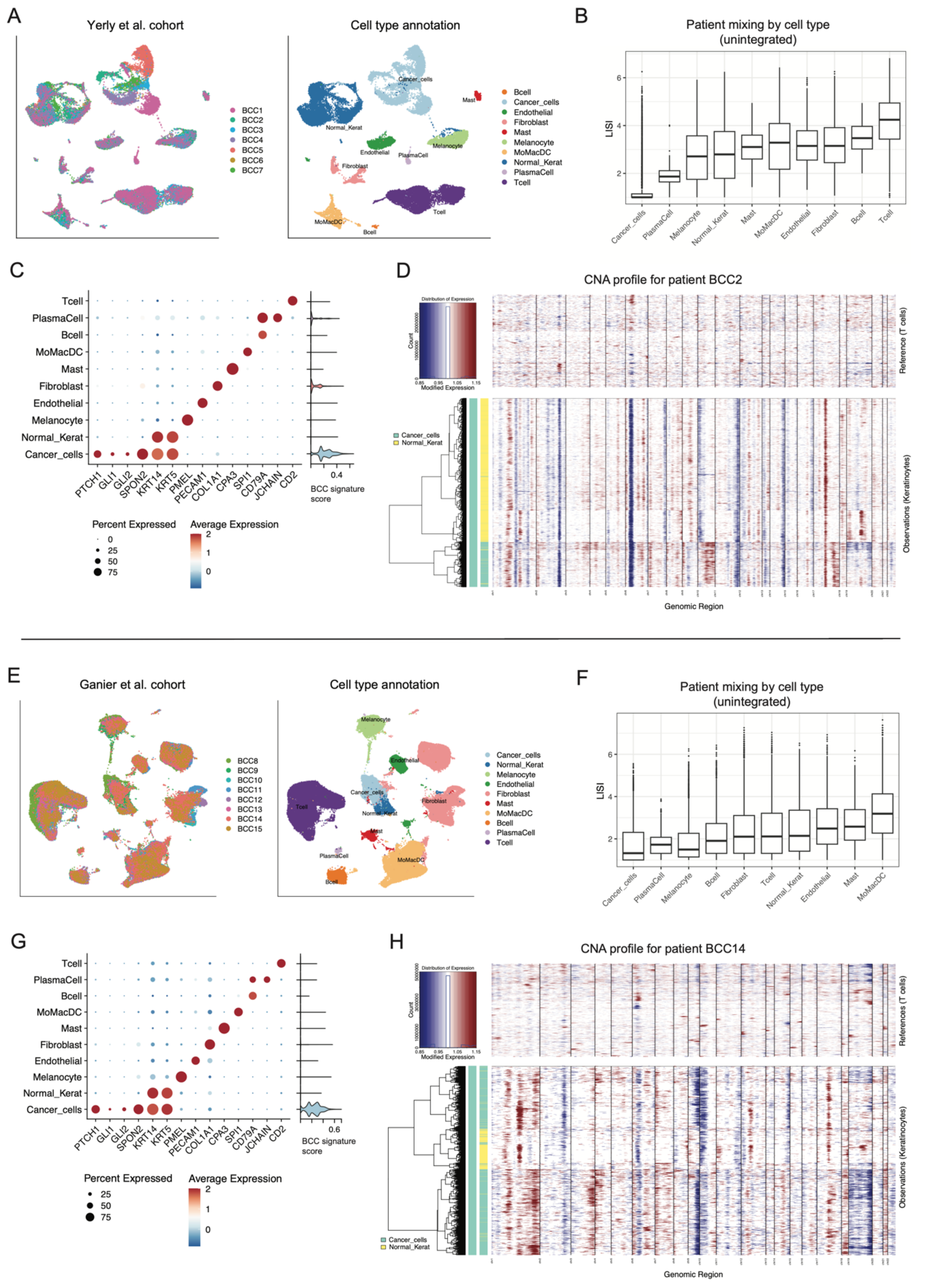


**Figure S1. Identification of cancer cells from scRNA-seq of two human BCC cohorts. A)** UMAP embeddings by patient and by cell type annotation from the Yerly et al. cohort. Broad annotation was obtained by applying the scGate automated tool and refined by clustering analysis. Identification of cancer cells was performed using three criteria (patient mixing, signature scoring, and copy-number variation), outlined respectively in panels B, C and D. **B)** Local Inverse Simpson Index (LISI) by cell type, quantifying the number of patients having cells in any given neighborhood of cells. Cancer cells are expected to have more patient-specific transcriptomes than other cell types, and primarily cluster together with cells from the same patient. **C)** Scaled average expression for marker genes of major TME cell types, and UCell score for a BCC signature defined by Yerly et al. (2022). **D)** Copy-number variation (CNV) profile for a representative patient, confirming that normal keratinocytes and cancer cells primarily cluster together in terms of inferred CNVs. **E)** UMAP embeddings by patient and by cell type annotation from the Ganier et al. cohort. Broad annotation was obtained by applying the scGate automated tool and refined by clustering analysis. Identification of cancer cells was performed using three criteria (patient mixing, signature scoring, and copy-number variation), outlined respectively in panels F, G and H. **F)** Local Inverse Simpson Index (LISI) by cell type, quantifying the number of patients having cells in any given neighborhood of cells. Cancer cells are expected to have more patient-specific transcriptomes than other cell types, and primarily cluster together with cells from the same patient. **G)** Scaled average expression for marker genes of major TME cell types, and UCell score for a BCC signature defined by Yerly et al. (2022). **H)** Copy-number variation (CNV) profile for a representative patient, confirming that normal keratinocytes and cancer cells primarily cluster together in terms of inferred CNVs.

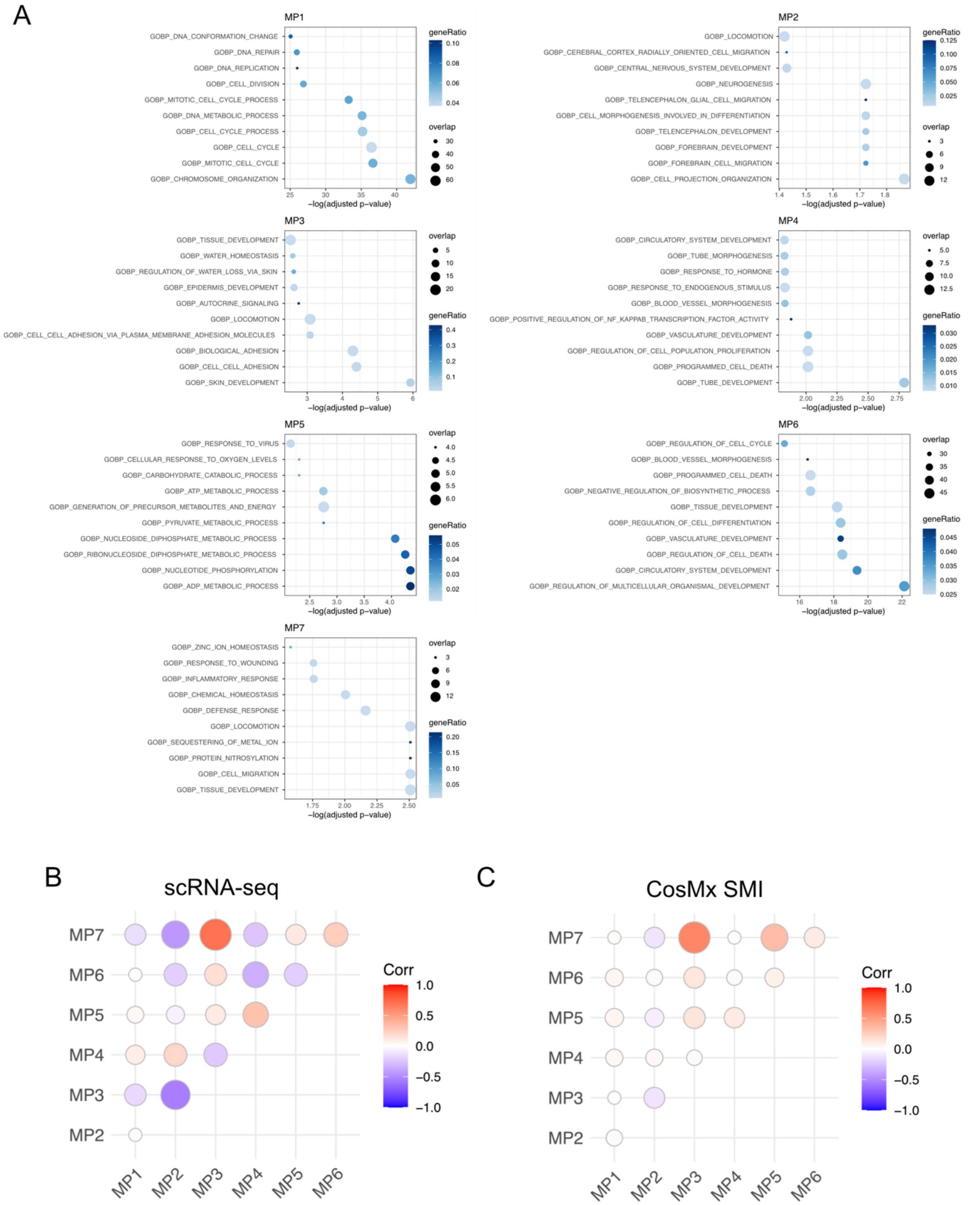


**Figure S2. Characterization of the meta-programs and their interrelation. A)** Gene set enrichment analysis (GSEA) between BCC meta-programs and Gene Ontology (GO) gene sets from MSigDB. Dot size is proportional to the number of genes shared between a MP-GO term pair, dot color represents the ratio between the number of shared genes over the total number of genes in the GO signature. **B)** Pairwise correlation coefficient between MP signature scores in cancer cells from scRNA-seq data (11 samples). **C)** Pairwise correlation coefficient between MP signature scores in cancer cells from CosMx SMI *in situ* transcripts.

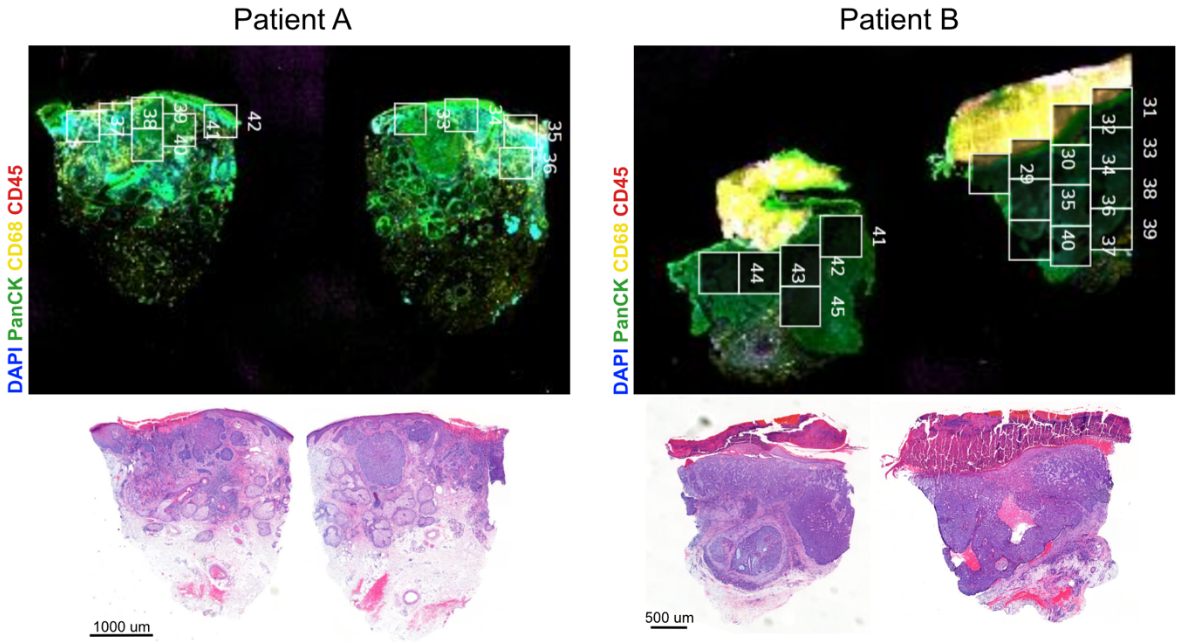


**Figure S3. Overview of tumor samples and Fields-of-View (FOVs) selection for CosMx SMI.** For Patient A & Patient B processed with the Cosmx SMI: (Top panels) Overview immunofluorescence image colored by DAPI (blue), pan-cytokeratin (green), CD68 (yellow) and CD45 (red), and annotated for FOVs selection. (Bottom panels) Overview morphological staining (H&E) from an adjacent slide.

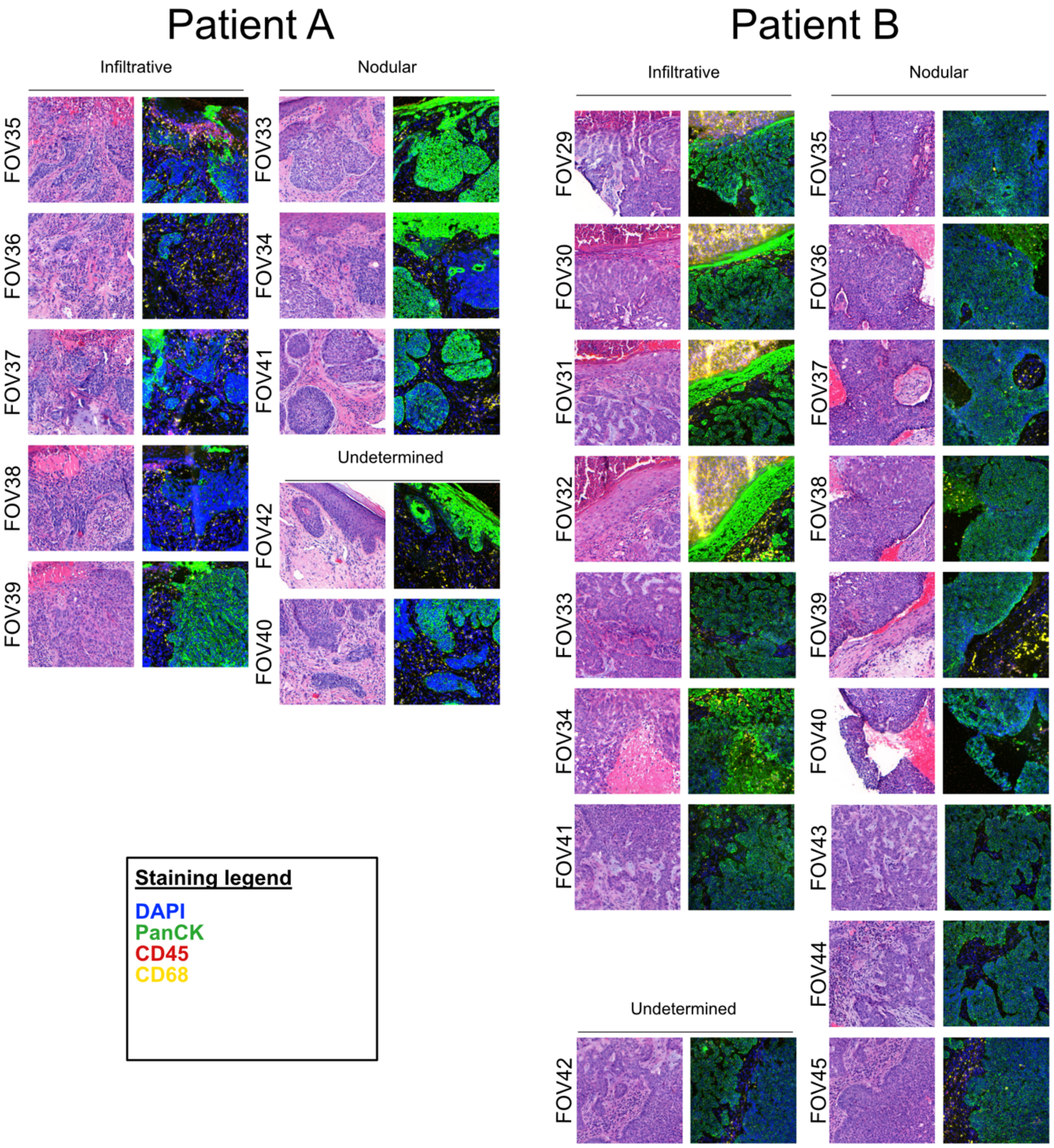


**Figure S4. Detailed H&E and immunofluorescence for individual FOVs in CosMx SMI.** Morphological staining (H&E) and immunofluorescent image of individual FOVs colored by DAPI (blue), pan-cytokeratin (green), CD68 (yellow) and CD45 (red), and annotated for infiltrative, nodular or undetermined morphology (patient A and B).

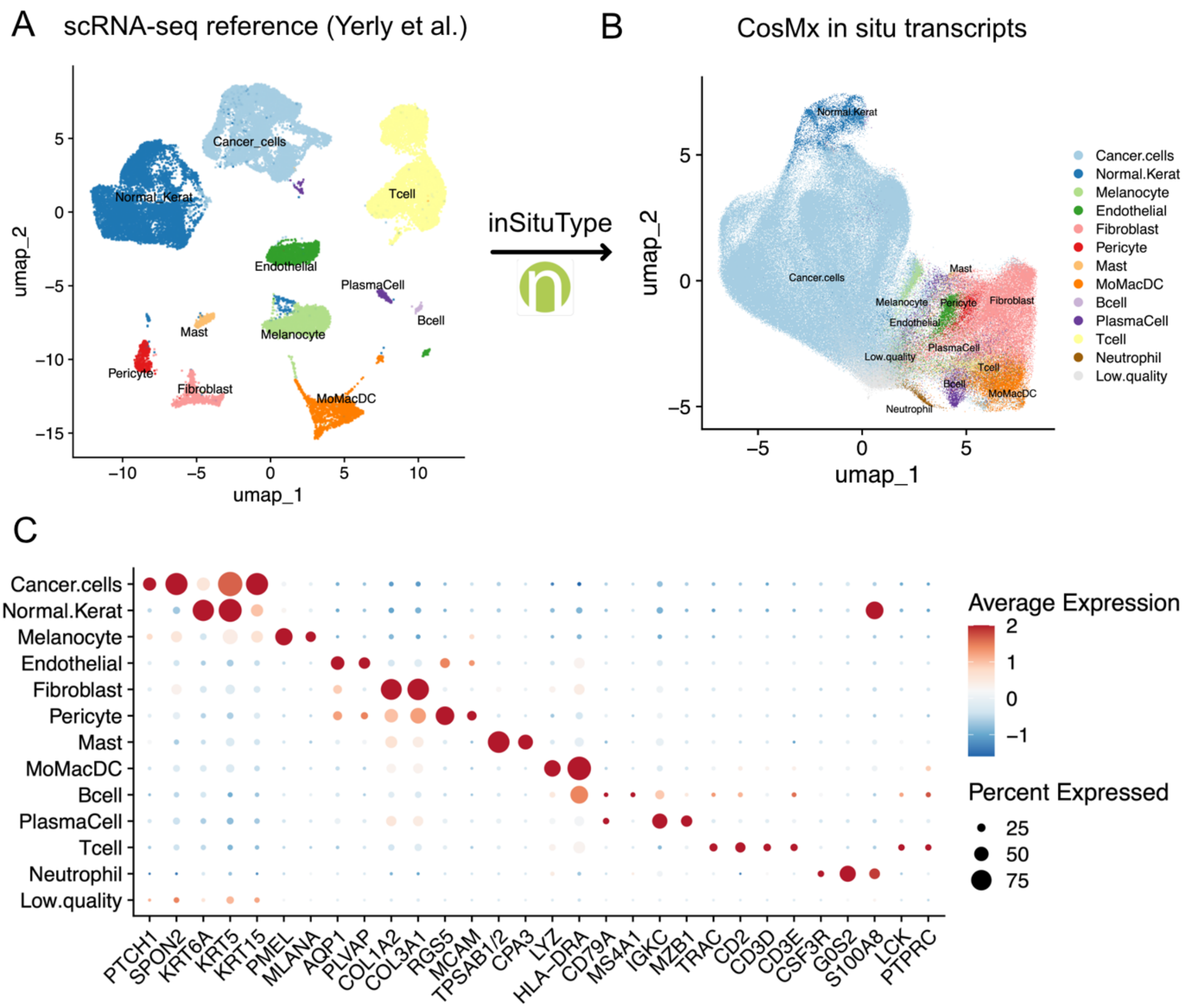


**Figure S5. Prediction of cell types in CosMx SMI *in situ* transcripts. A)** UMAP embeddings for scRNA-seq samples from Yerly et al. (2022) **B)** Based on the reference expression profiles of the data in A) – plus the expression profile for neutrophils derived from Zilionis et al. (2019) – the inSituType algorithm was applied to transfer labels to the *in situ* transcripts of 4 patients sequenced with CosMx SMI technology. **C)** Normalized average expression (color scale) and percent of expressing cells (dot size) for a panel of marker genes, for each of the cell types defined in the CosMx SMI dataset.

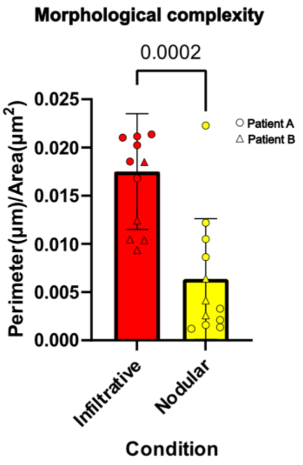


**Figure S6. Morphological complexity of BCCs measured as perimeter/area ratio, in infiltrative vs. nodular Fields-of-View (FOVs).** Morphological complexity measured as perimeter/area ratio on the composite images from Cosmx SMI based on the DAPI (blue) and pan-cytokeratin (green) stainings for FOVs of 2 patients previously annotated as Nodular or Infiltrative. P-value: unpaired t-test.

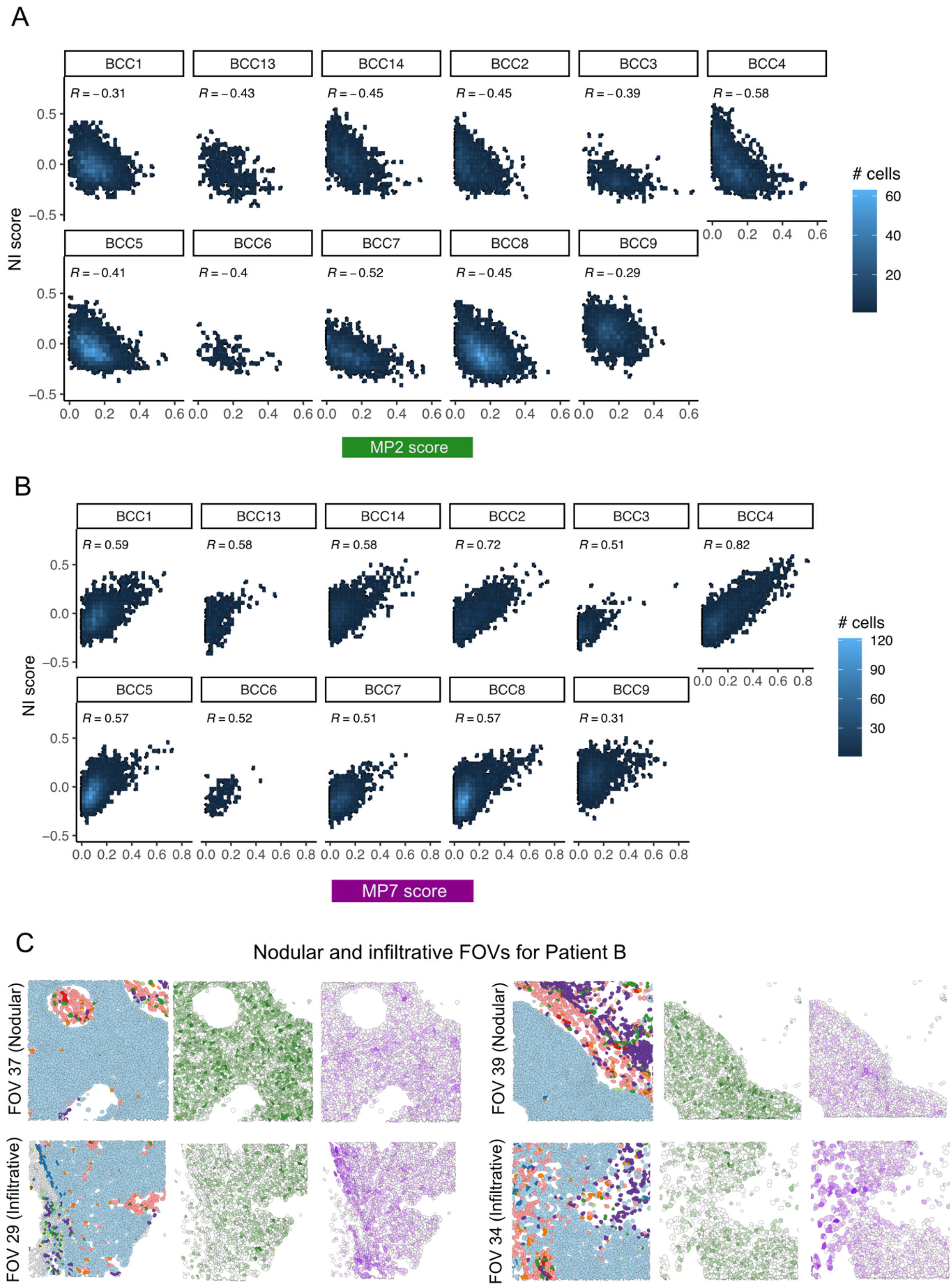


**Figure S7. Correlation between MPs and nodular-to-infiltrative score (NI score). A)** Density plots for the correlation of MP2 with the NI score (see Methods) in scRNA-seq from 11 BCC patients. **B)** Density plots for the correlation of MP7 with the NI score in scRNA-seq from 11 BCC patients. The correlation coefficient is reported on top of each plot; color scale indicates the density of cells. **C)** Predicted cell types and spatial distribution of MP2 and MP7 in four representative FOVs from Patient B (two nodular, two infiltrative).

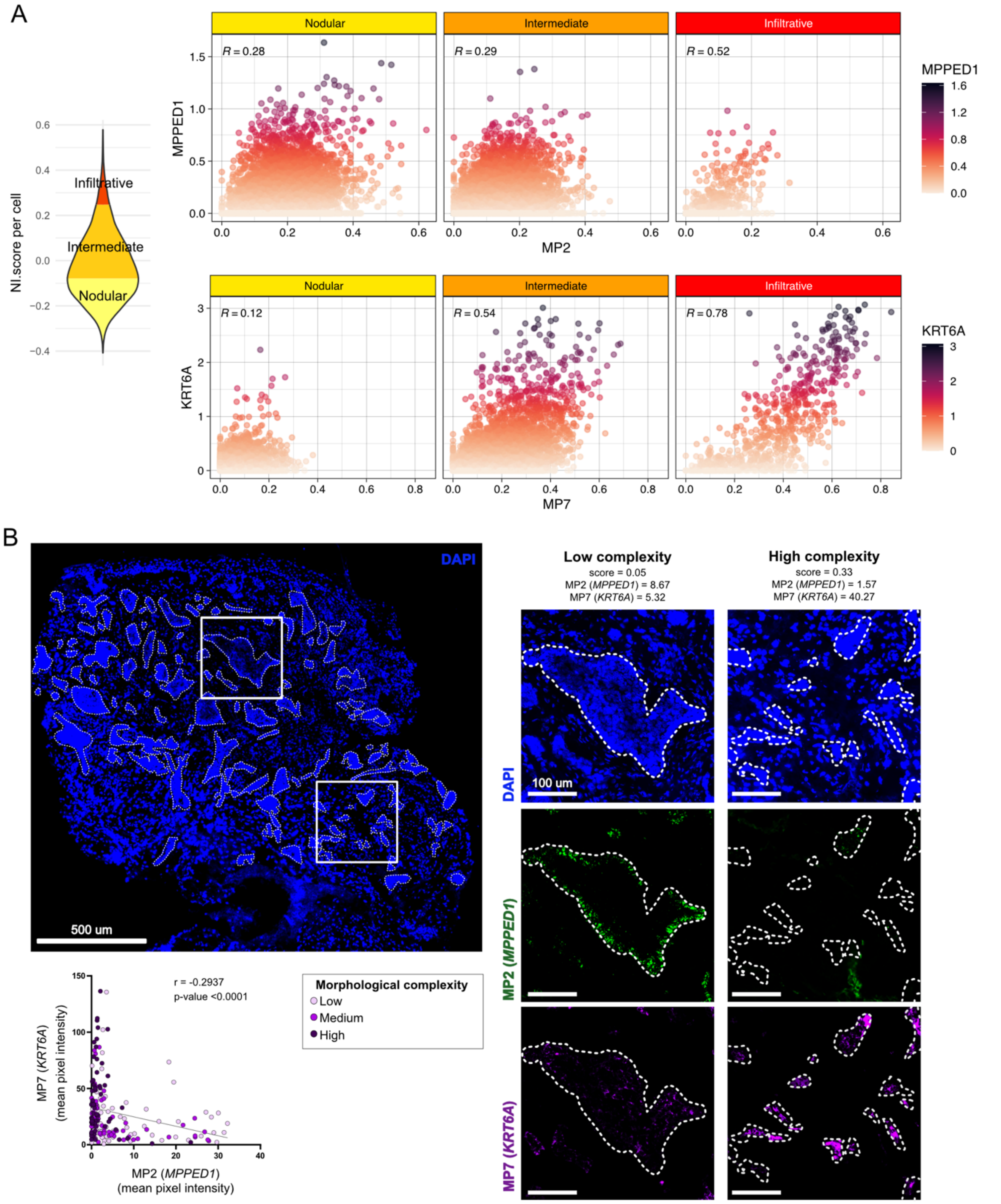


**Figure S8. Validation of MP2 and MP7 markers. A)** Expression of *MPPED1* vs. MP2 signature score for cells with nodular, intermediate and infiltrative transcriptomics profiles; and expression of *KRT6A* vs. MP7 signature score for cells with nodular, intermediate and infiltrative transcriptomics profiles. Cells are classified in these three categories by binning the observed NI score range into three equal intervals. **B)** Overview immunofluorescent image stained with DAPI of a human BCC with high intratumoral morphological heterogeneity. White dotted lines indicate tumor areas measured for morphological complexity (perimeter/area) and mean pixel intensity of MP7 (*KRT6A*) and MP2 (*MPPED1*) markers (top left panel). Correlation analysis between MP7 (*KRT6A*) and MP2 (*MPPED1*) markers (bottom left panel). Each dot is colored according to its morphological complexity score (low, medium, high). Illustrative images of low and high complexity tumor areas at higher magnification colored with DAPI (blue), *MPPED1* (green) and *KRT6A* (purple) probes for RNA FISH (right panels). P-value and r: Spearman correlation.

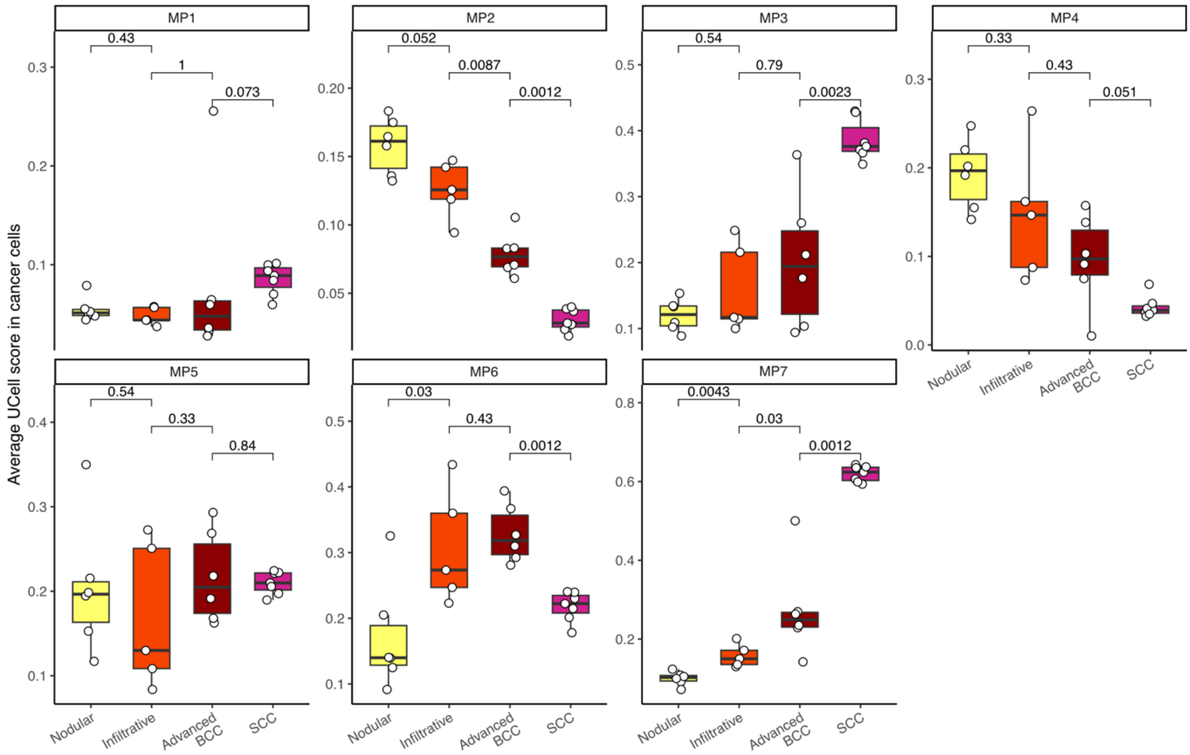


**Figure S9. Meta-program scores in skin cancers of increasing aggressiveness.** Signature scores for MP1-MP7 in scRNA-seq data of nodular and infiltrative BCCs (Yerly et al. 2022); advanced BCCs (Yost et al. 2019); and cSCCs (Ji et al. 2020). Each point represents the average MP score in cancer cells of one sample. cSCC: cutaneous squamous cell carcinoma.

**A**

| Sample | Age | Gender | Location | Treatment | Naive | Resistant |
|---|---|---|---|---|---|---|
| Patient 1 | 60 | F | Thorax | Vismodegib | 1 | |
| Patient 2 | 30 | F | Thorax | Sonidegib | 1 | |
| Patient 3 | 81 | F | Back | Sonidegib | 1 | |
| Patient 4 | 83 | M | Left temple | Vismodegib | 2 | 1 (7 months) |
| Patient 5 | 68 | M | Left thorax | Sonidegib and Vismodeg | | 1 (18 months) |
| Patient 6 | 66 | M | Right leg | Vismodegib | 1 | 1 (6 weeks)<br>1 (8 months) |
| Patient 7 | 74 | F | Periocular (Left eye) | Sonidegib | 1 | |

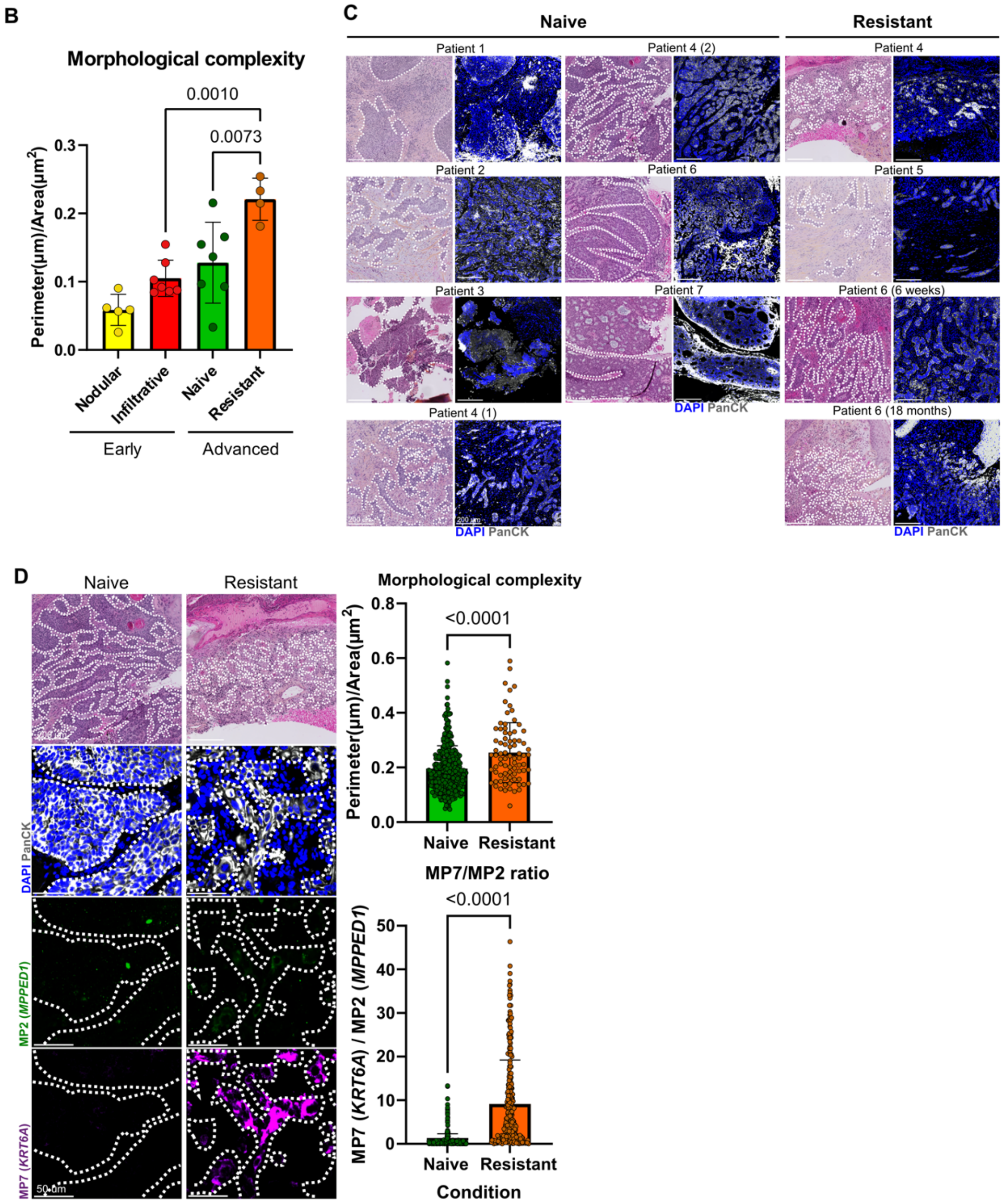


**Figure S10. Morphological features of advanced, therapy-resistant BCCs. A)** Metadata table summarizing advanced BCCs patient's characteristics. **B)** Morphological complexity measured as perimeter/area ratio in early tumors previously annotated as nodular or infiltrative BCCs by dermatopathologists and in advanced BCCs naive or resistant to HHI treatment. **C)** Representative H&E and immunofluorescent images stained with DAPI (blue) and pan-cytokeratin (grey) of individual advanced BCC samples classified as naive or resistant to treatment. **D)** Patient-matched treatment-naive and treatment-resistant samples shown for representative H&E and immunofluorescent images (left panels), and measured for morphological complexity (measured by perimeter/area ratio; top right panel) and MP7/MP2 ratio (assessed by the ratio of mean pixel intensity of MP7 marker *KRT6A* over mean pixel intensity of MP2 marker *MPPED1;* bottom right panel). P-values: unpaired Student's *t* test.

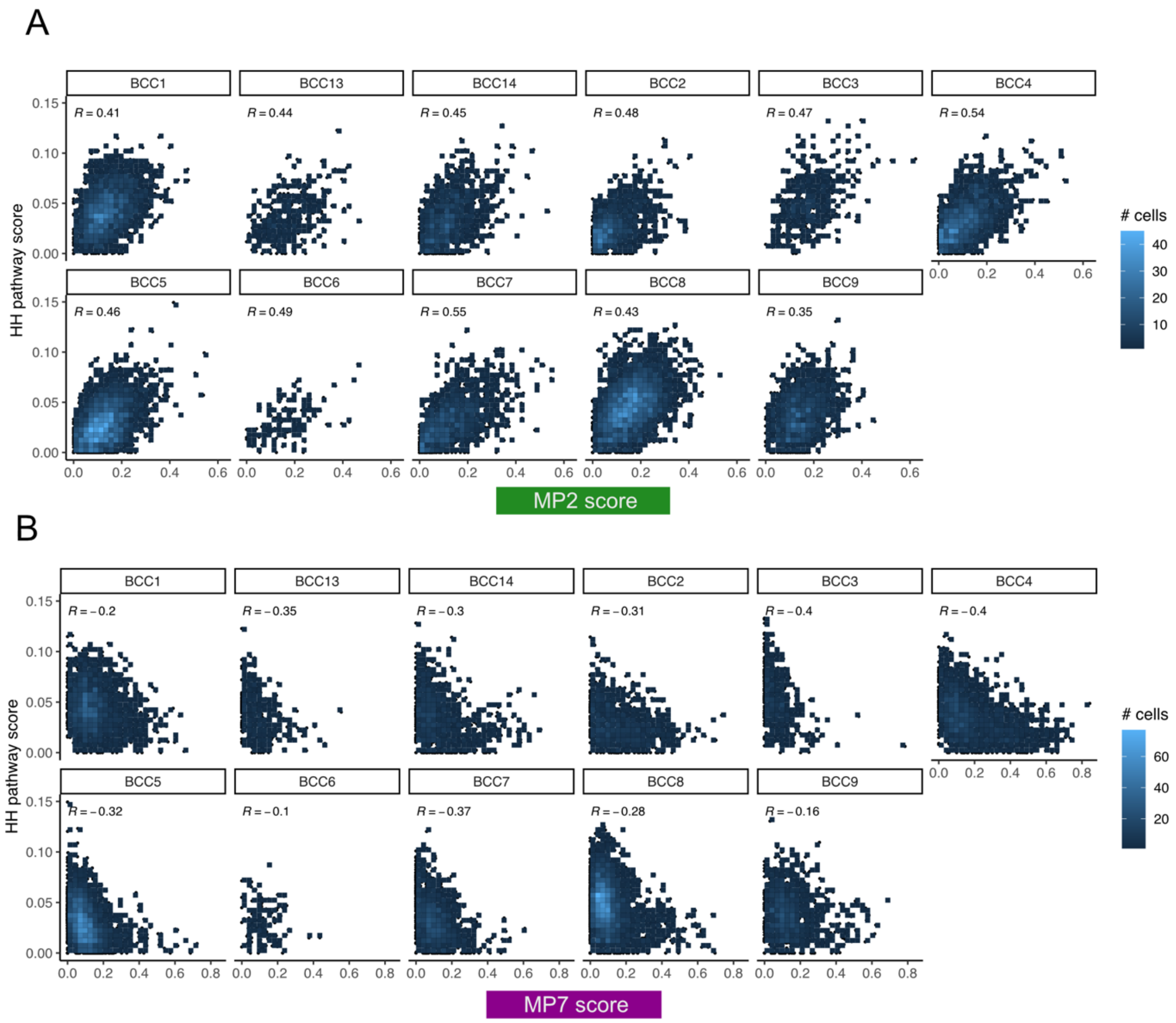


**Figure S11. Correlation between MPs and hedgehog (HH) pathway score. A)** Density plots for the correlation of MP2 with the HH pathway score (from KEGG database) in scRNA-seq from 11 BCC patients. **B)** Density plots for the correlation of MP7 with the HH pathway score in scRNA-seq from 11 BCC patients. The correlation coefficient is reported on top of each plot; color scale indicates the density of cells.

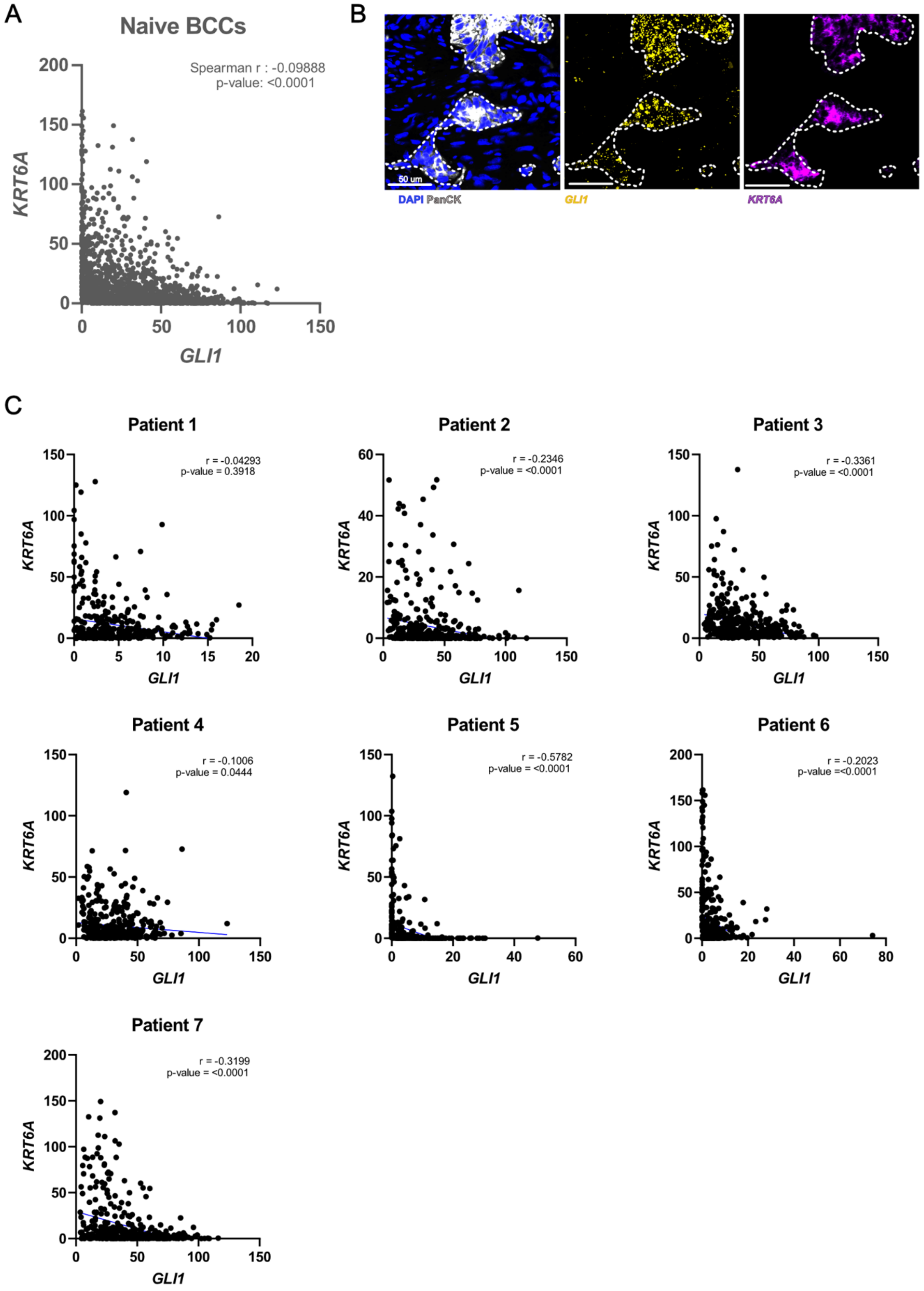


**Figure S12. Relation between MP7 and HH pathway markers. A)** Correlation analysis of mean pixel intensity of MP7 marker K*RT6A* and Hedgehog pathway activity G*LI1* at single-cell level across 7 human BCCs. **B)** Representative immunofluorescent images stained with DAPI (blue), pan-cytokeratin (grey), *GLI1* (yellow) and *KRT6A* (purple) probes for RNA FISH. **C)** Correlation analysis of mean pixel intensity of MP7 marker *KRT6A* and Hedgehog pathway activity marker *GLI1* at single-cell level for individual patients. R and p-values: Spearman correlation.

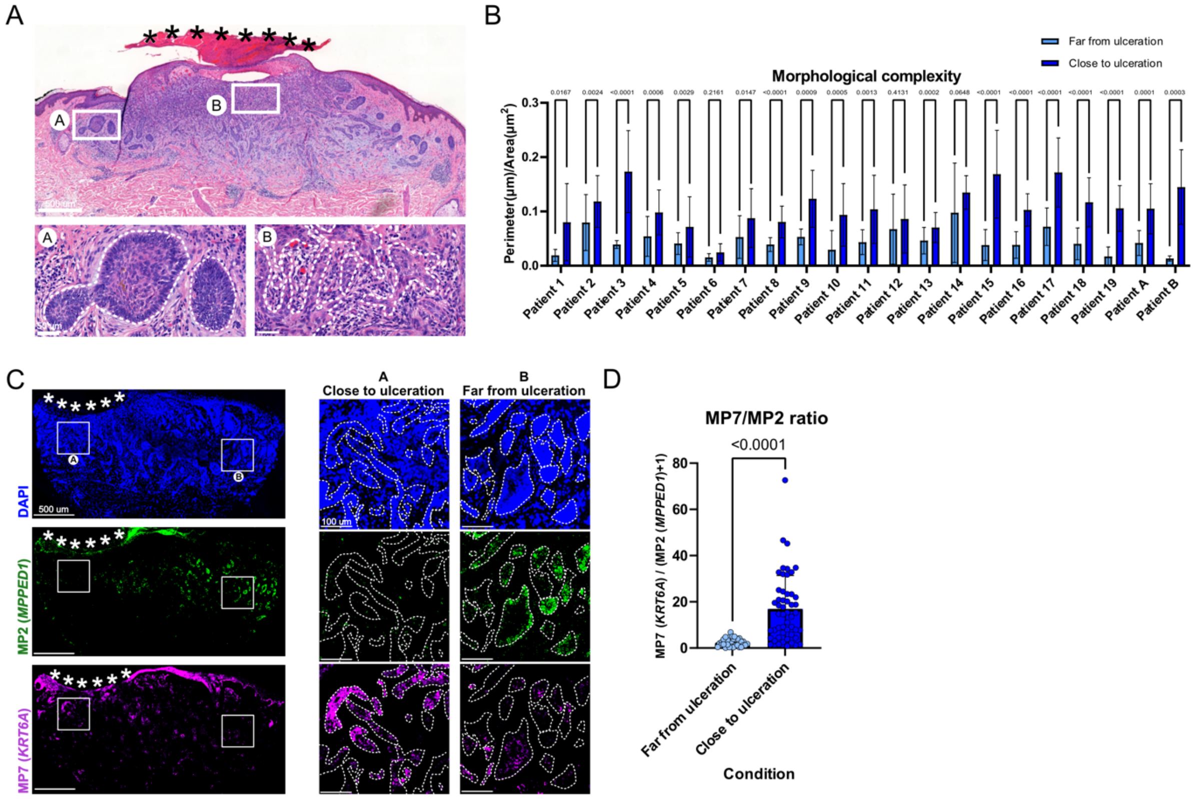


**Figure S13. Morphological complexity in ulcerated BCCs. A)** Representative scan of H&E images of an ulcerated BCC sample in which (A) represents an area "far from ulceration" and (B) an area "close to ulceration". White dotted lines depict the basement membrane zone (BMZ). **B)** Mean tumor complexity in area far and close to ulceration (N≥6 tumor structures per condition of 21 different BCC samples. Horizontal bars indicate the mean ± SD, p-values were calculated by unpaired two-sided Student's *t* test. **C)** Representative scan of immunofluorescent images of an ulcerated BCC sample colored with DAPI (blue), *MPPED1* (green) and *KRT6A* (purple) probes for RNA FISH. (A) represents an area "far from ulceration" and (B) an area "close to ulceration. White dotted lines depict the BMZ. D) MP7/MP2 ratio assessed by the ratio of mean pixel intensity of MP7 marker *KRT6A* over mean pixel intensity of MP2 marker *MPPED1.* P-value: unpaired Student's *t* test.

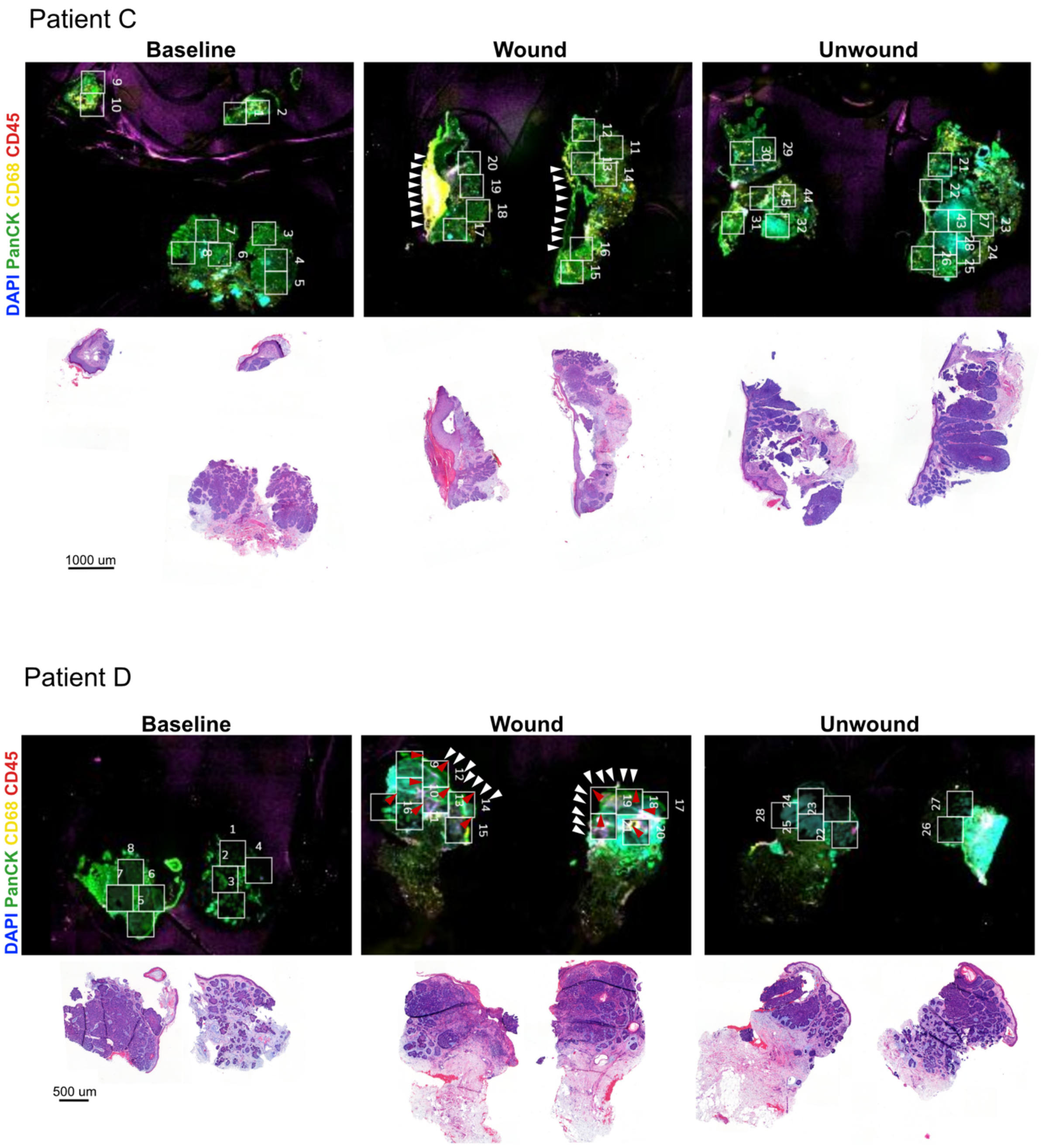


**Figure S14. Overview of tumor samples and Fields-of-View (FOVs) selection for CosMx SMI.** For Patient C & Patient D processed with the Cosmx SMI: (Top panels) Overview immunofluorescence image colored by DAPI (blue), pan-cytokeratin (green), CD68 (yellow) and CD45 (red), and annotated for FOVs selection. (Bottom panels) Overview morphological staining (H&E) from an adjacent slide.

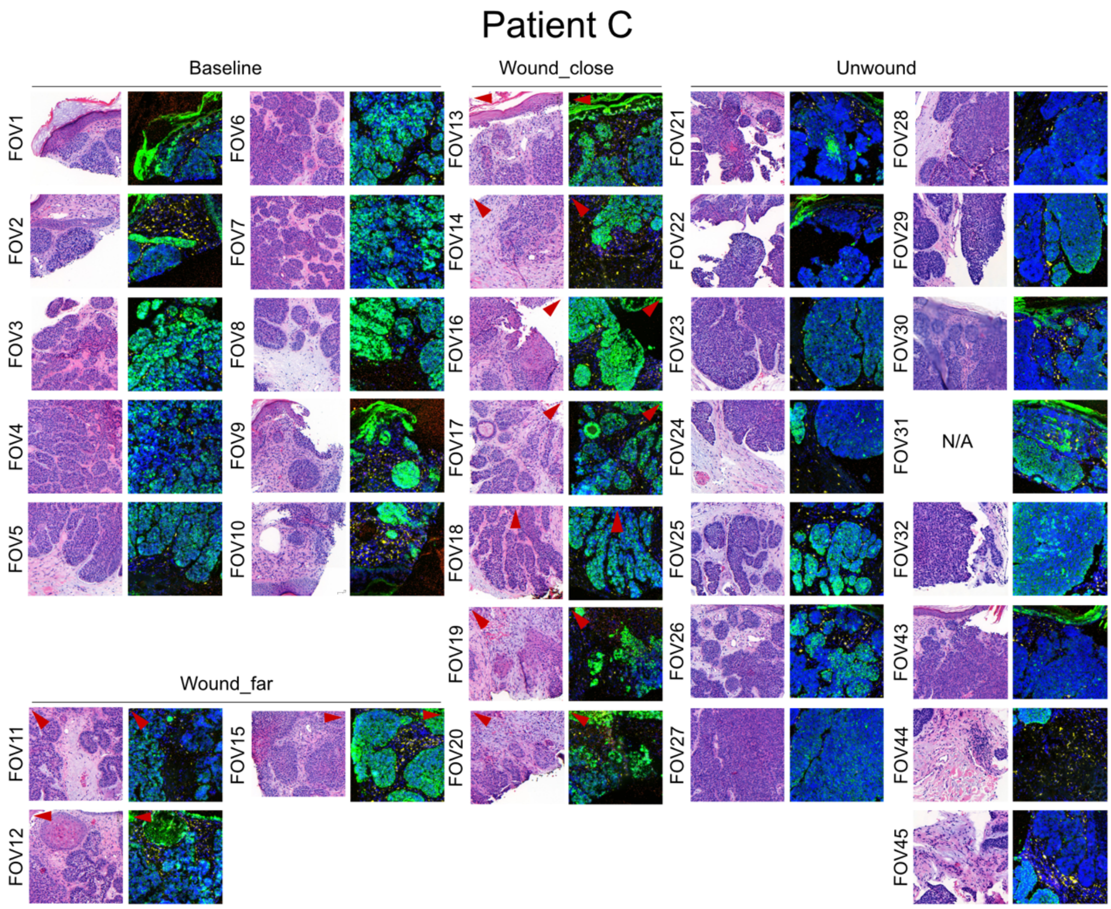
Patient C
Baseline
Wound_close
Unwound
Wound_far
FOV1
FOV2
FOV3
FOV4
FOV5
FOV6
FOV7
FOV8
FOV9
FOV10
FOV11
FOV12
FOV13
FOV14
FOV15
FOV16
FOV17
FOV18
FOV19
FOV20
FOV21
FOV22
FOV23
FOV24
FOV25
FOV26
FOV27
FOV28
FOV29
FOV30
FOV31
N/A
FOV32
FOV43
FOV44
FOV45

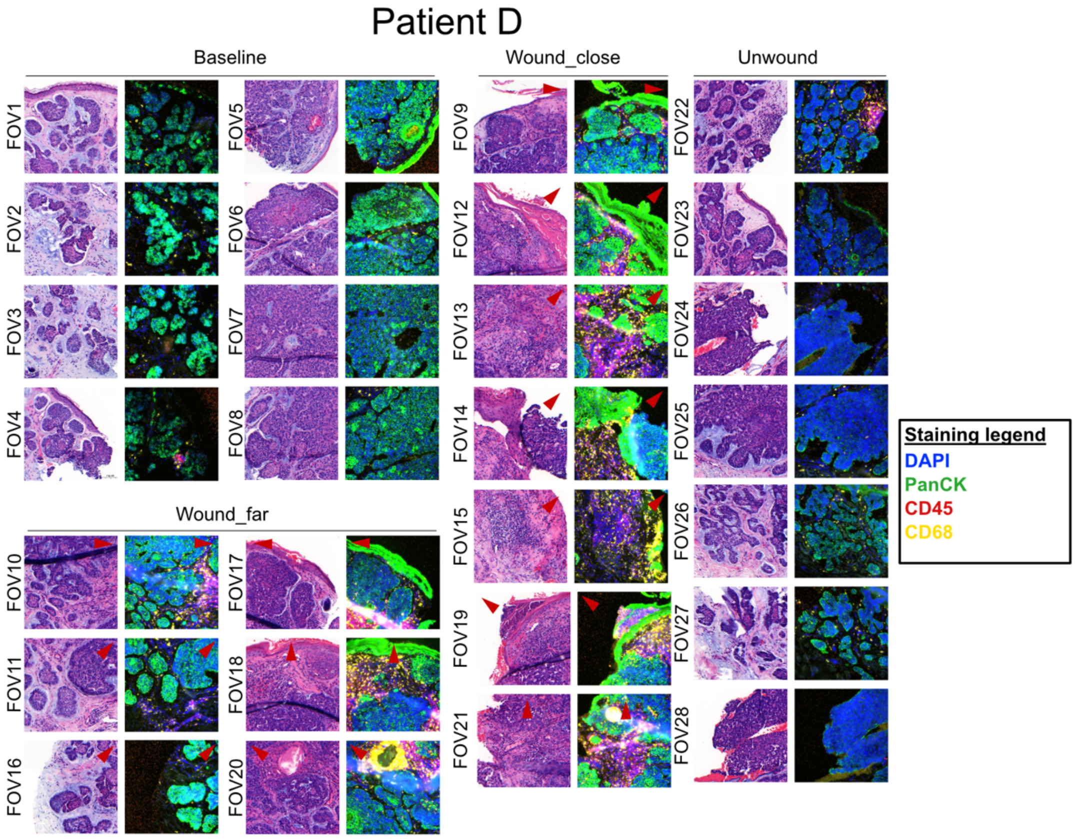
Patient D
Baseline
Wound_close
Unwound
Wound_far
FOV1
FOV2
FOV3
FOV4
FOV5
FOV6
FOV7
FOV8
FOV9
FOV10
FOV11
FOV12
FOV13
FOV14
FOV15
FOV16
FOV17
FOV18
FOV19
FOV20
FOV21
FOV22
FOV23
FOV24
FOV25
FOV26
FOV27
FOV28
Staining legend
DAPI
PanCK
CD45
CD68

**Figure S15. Detailed H&E and immunofluorescence for individual FOVs in CosMx SMI.** Morphological staining (H&E) and immunofluorescent image of individual FOVs colored by DAPI (blue), pan-cytokeratin (green), CD68 (yellow) and CD45 (red), and annotated for baseline, wound_far_wound_close and unwound condition (patient C and D). Red arrow heads indicate direction to the wound. N/A: Not available.

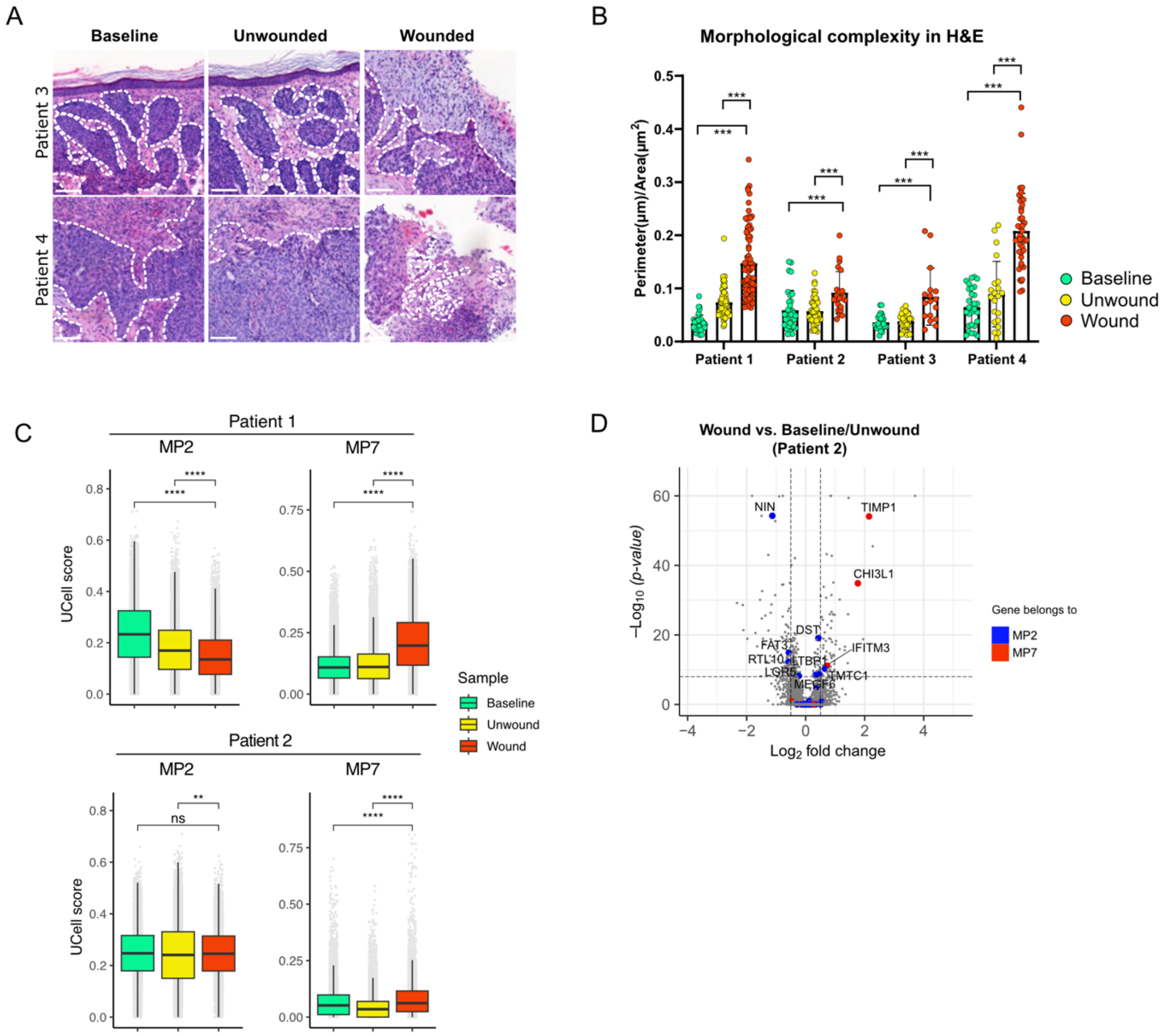


**Figure S16. Morphological and transcriptional changes upon BCC wounding. A)** H&E staining for patients 3 and 4 at first biopsy (baseline), at day 7 far from the first biopsy (unwounded), and at day 7 on the same site as the first biopsy (wounded). **B)** Morphological complexity, measured as perimeter/area ratio, for multiple areas of four BCC patients, in the three indicated conditions. **C)** UCell signature score for MP2 and MP7 meta-programs in scRNA-seq from two BCC patients, in the three indicated conditions. **D)** Differentially expressed genes between Wound vs. Baseline/Unwound samples for Patient 2 (data for the other patient in Figure 4). Genes contained in MP2 and MP7 are highlighted blue and red respectively.

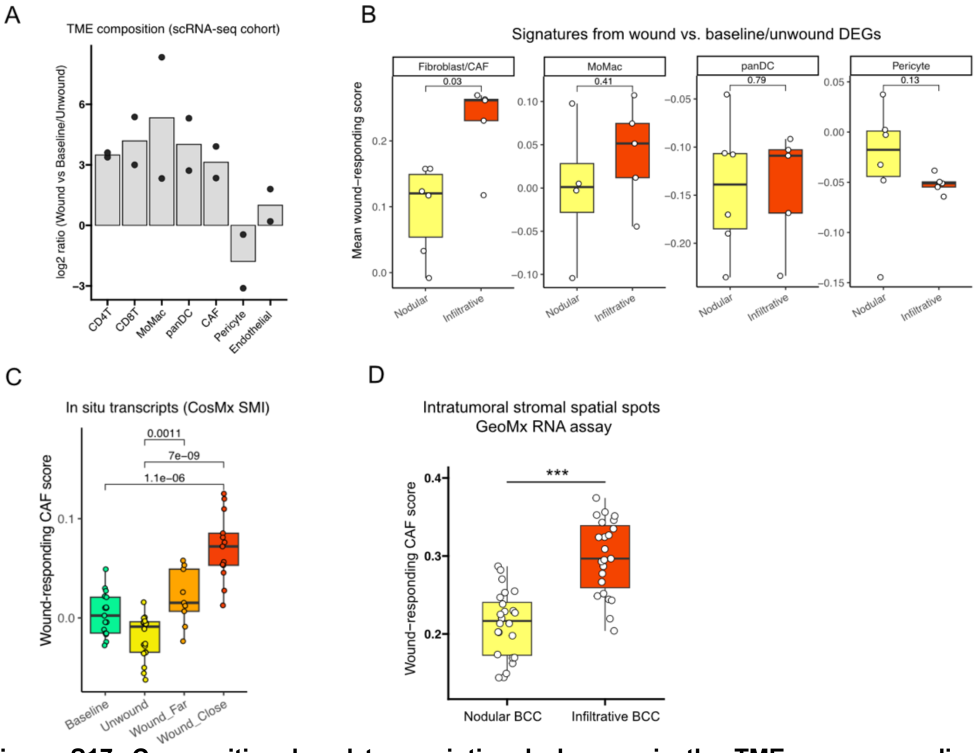


**Figure S17. Compositional and transcriptional changes in the TME upon wounding. A)** Compositional changes in the TME upon wounding, measured in a scRNA-seq cohort. For each cell type, the enrichment score is calculated as the log-ratio of the cell type frequency in wounded vs. baseline/unwounded samples (N=2). **B)** Signature scores for the gene sets derived by differential expression analysis in wounded vs. baseline samples, for fibroblasts (CAF), monocyte/macrophages (MoMac), all dendritic cells (panDC) and pericytes, evaluated in 11 nodular and infiltrative BCC scRNA-seq samples. **C)** Wound-responding CAF score in a spatial transcriptomics dataset, averaged by FOVs from baseline, unwounded and wounded samples (far or close to the wound). **D)** Wound-responding CAF score calculated in intratumoral stromal spatial spots from a GeoMx RNA experiment (Yerly et al. 2022), where BCC areas were previously annotated as nodular or infiltrative. p-values: Wilcoxon rank-sum test.
